# Pervasive and recurrent hybridisation prevents inbreeding depression in Europe’s most threatened seabird

**DOI:** 10.1101/2024.10.24.619781

**Authors:** Guillem Izquierdo-Arànega, Cristian Cuevas-Caballé, Francesco Giannelli, Josephine Rosanna Paris, Karen Bourgeois, Emiliano Trucchi, Jacob González-Solís, Marta Riutort, Joan Ferrer Obiol, Julio Rozas

**Affiliations:** Departament de Biologia Evolutiva, Ecologia i Ciències Ambientals, Universitat de Barcelona, Barcelona, Spain; Institut de Recerca de la Biodiversitat, Universitat de Barcelona, Barcelona, Spain; Departament de Genètica, Microbiologia i Estadística, Universitat de Barcelona, Barcelona, Spain; Department of Life and Environmental Sciences, Marche Polytechnic University, Ancona, Italy; Aix Marseille Université, CNRS, IRD, Avignon Université, Institut Méditerranéen de Biodiversité et d’Ecologie marine et continentale, Bât. Villemin, Technopôle Arbois-Méditerranée, UMR IMBE, Aix-en-Provence, France; Dipartimento di Scienze e Politiche Ambientali, Università degli Studi di Milano, Milano, Italy

## Abstract

Hybridisation is a double-edged sword: while it can erode distinct evolutionary lineages, it can also introduce genetic diversity and adaptive potential into dwindling populations. In the Critically Endangered Balearic shearwater (*Puffinus mauretanicus*), this dilemma is exacerbated by a limited understanding of the extent and consequences of hybridisation with the Yelkouan shearwater (*P. yelkouan*). This knowledge gap has limited the scope of science-based conservation strategies to avoid the Balearic shearwater’s imminent extinction. Here, we investigate shearwater hybridisation dynamics and their effect on genome-wide diversity in the Balearic shearwater. Divergence dating, demographic modelling and admixture analyses suggest that these two poorly-differentiated shearwater lineages experienced recurrent episodes of divergence and widespread hybridisation during glacial cycles. Selection scans reveal a 500 kb region hosting an adaptive haplotype that potentially underpins interspecific differences in migratory behaviour, and which has been repeatedly introgressed between taxa. Moreover, we show that interspecific gene flow has prevented increases in homozygosity and genetic load, and through forward simulations we illustrate how it can enhance the Balearic shearwater’s resilience to future population bottlenecks. Our findings illustrate how introgression can be crucial for maintaining genetic diversity in threatened taxa, and highlight the need for considering the protection of hybridisation in conservation plans.

## INTRODUCTION

Hybridisation has traditionally been viewed as a problem for species conservation. This process can lead to the merging of distinct lineages and can threaten endangered taxa by introducing maladaptive genetic variation^1,2^. Hence, it has been argued that hybridisation can ultimately increase the extinction risk of a species^1,2^. However, this view is being increasingly challenged by research that highlights the role of hybridisation in boosting the evolutionary potential of taxa, either through the introduction of adaptive mutations or through the generation of novel haplotypes^3–5^. Small and depleted populations may particularly benefit from this process. Inbreeding makes these populations especially vulnerable to extinction by increasing homozygosity and reducing genetic diversity, which can lead to the loss of both individual fitness and adaptive potential^6–9^. Hybridisation can counteract these effects and rescue a population from falling into the extinction vortex through natural genetic rescue^3–5,10^. This shift in mindset has led to calls for the inclusion of hybrid populations in conservation and management plans^5,10^.

The conservation outlook for the Balearic shearwater (*Puffinus mauretanicus*) is dire: this Critically Endangered seabird, endemic to the Balearic Islands (∼3,000 breeding pairs^11^), has declined dramatically over recent decades (∼14% annual decline^12^) and a recent viability analysis found that the species could face extinction by 2070^12^. This precipitous decline has been driven by longline bycatch and predation by alien species^11^. The effect of these declines on the species’ inbreeding risk remains unknown. Yet, the Balearic shearwater might be facing an additional threat due to hybridisation with its sister species, the Yelkouan shearwater (*P. yelkouan*)^13^. Although also threatened, the Yelkouan shearwater has a much wider distribution, breeding throughout the rest of the Mediterranean Sea. The two taxa differ markedly in mtDNA^14,15^, plumage, size^16^ and migratory behavior^17^ (Fig. 1a), and are currently treated as separate species (although see Ferrer Obiol *et al.* (2023)^18^). Nevertheless, interspecific hybridisation is suspected to occur in Menorca, the easternmost Balearic Island, which is inhabited by a morphologically and ecologically intermediate population that shares mitochondrial haplotypes from both taxa^13,14,17^.

**Fig. 1:**
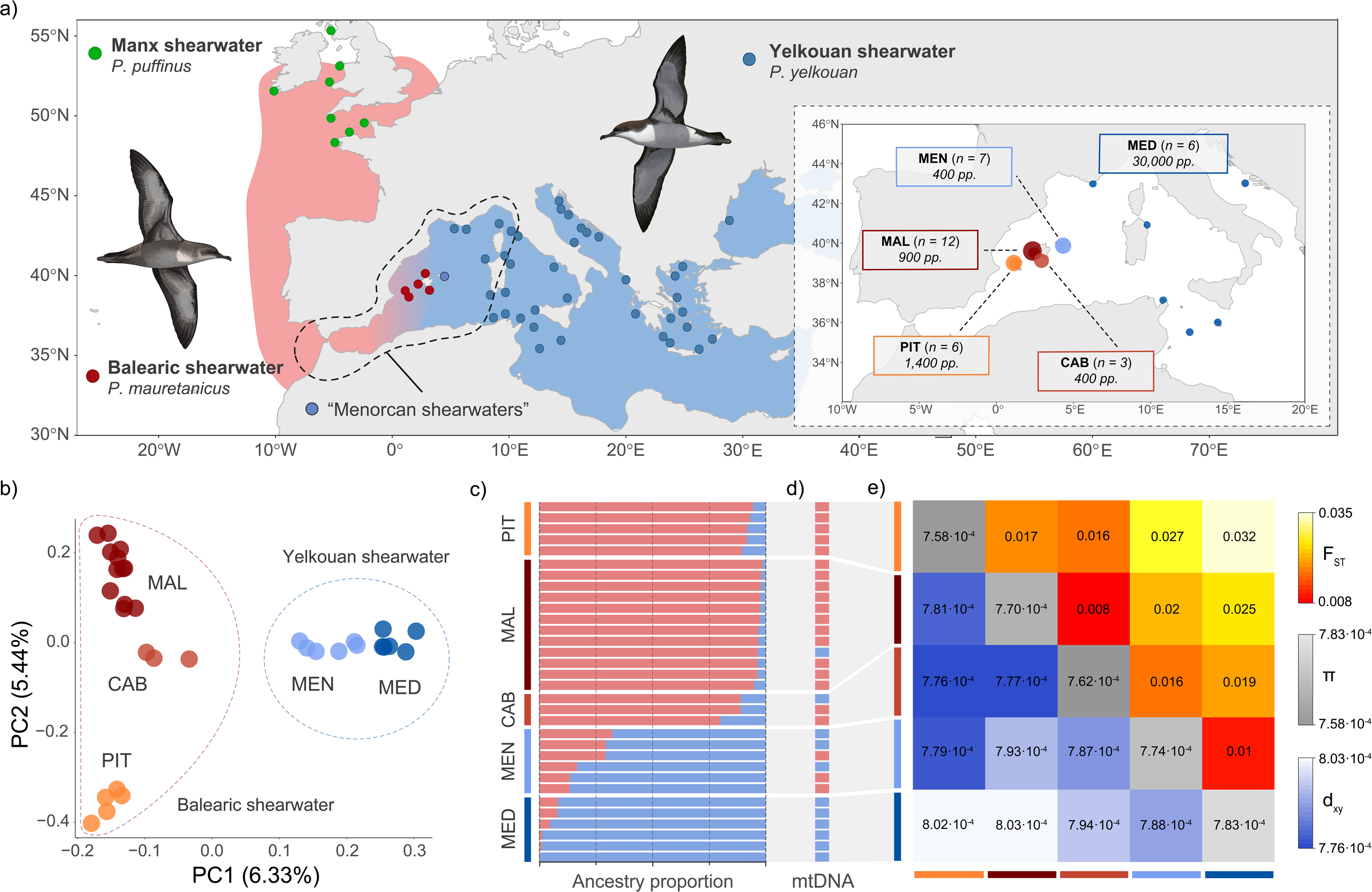
Distribution and population structure across Mediterranean *Puffinus* shearwaters. **a,** Non-breeding distribution (shaded areas) and breeding colonies (dots) of the *Puffinus* shearwater taxa included in this study (adapted from Austin *et al.* (2019)^17^ and Harrison *et al.* (2020)^100^). Inset shows the sampling locations with sample sizes (in parentheses) and estimated census sizes in breeding pairs (pp.). **b,** PCA of the sampled Mediterranean *Puffinus* based on 1,304,832 autosomal SNPs. **c,** Struct-f4 ancestry component profiles of the sampled *Puffinus* assuming *K = 2* genetic ancestries. **d,** Mitochondrial haplogroups assuming *K = 2* mitochondrial clades. **e,** Colour-coded heatmaps according to pairwise differentiation (*F_ST_*; above diagonal), absolute divergence (*d_xy_*; below diagonal) and nucleotide diversity (*π*; diagonal) across populations. Illustrations of Balearic (bottom left) and Yelkouan shearwaters (top right) by Martí Franch©.

Consequently, there is an urgent need to: (1) reconstruct the evolutionary history of *Puffinus* shearwater populations in the Mediterranean; (2) estimate the extent of hybridisation between them; and (3) evaluate the potential effects of inbreeding resulting from current population declines. Here, we use whole-genome resequencing data to address these questions and to estimate relevant evolutionary parameters crucial for conservation plans. Our findings reveal that recurrent bouts of pervasive gene flow during the Pleistocene interglacial periods have led to: (1) the discordance in coalescence times between mitochondrial and nuclear genomes, and (2) the exchange of adaptive alleles across taxa. Furthermore, we show that hybridisation has ensured the genetic viability of Balearic shearwaters and, via forward simulations, we provide compelling evidence of the conservation benefits of preserving genetic connectivity with the Yelkouan shearwater in the future.

## RESULTS

### Sampling, whole-genome resequencing and population structure

To investigate the evolutionary history of Mediterranean *Puffinus*, we generated whole-genome resequencing data for 36 individuals sampled across their breeding range. These included the three main extant populations of Balearic shearwater (Pitiüses (PIT, *n* = 6), Mallorca (MAL, *n* = 12) and Cabrera (CAB, *n* = 3)); the Menorcan shearwater population (MEN, *n* = 7); and six colonies of Yelkouan shearwater across the central Mediterranean Sea (MED, *n* = 6) (Fig. 1a). We also sequenced two individuals of their outgroup^18^, the Manx shearwater (*P. puffinus*) (Supplementary Table 1). Sequences were mapped to the Balearic shearwater reference genome and mitogenome^19^, resulting in an average sequence read depth of 8.31x (SD = 1.12) and 48.22x (SD = 19.61), respectively (Supplementary Table 2).

Population structure analyses using 1,304,832 genome-wide SNPs revealed a limited degree of genomic differentiation between Mediterranean *Puffinus* shearwater taxa. Principal component analysis (PCA) separated Balearic and Yelkouan shearwaters along PC1, yet the genomic variance explained (6.33%) was only marginally greater than the variance explained by intraspecific structure within Balearic shearwaters, as summarised along PC2 (5.44%; Fig. 1b). Individuals sampled from Menorca (MEN) clustered with the central Mediterranean (MED), suggesting that the former population belongs to Yelkouan shearwaters, unlike previously thought^14^ (Fig. 1b). Admixture analyses using Struct-f4^20^ for *K* = 2 populations delineated the same clusters as the PCA (Fig. 1c; Supplementary Fig. 1). However, several CAB and MEN individuals were inferred to share various proportions of ancestry from Balearic and Yelkouan shearwaters (Fig. 1c). Mitochondrial analysis identified two markedly divergent haplogroups, which separated most Balearic and Yelkouan shearwaters (Fig. 1d; Supplementary Fig. 2). Nevertheless, we found both mtDNA haplogroups within MAL, CAB and MEN (Fig. 1d; Supplementary Fig. 2). Interspecific pairwise *F_ST_* values were low (*F_ST_* = 0.016 - 0.032) and overlapped with intraspecific comparisons (*F_ST_* = 0.009 - 0.017) (Fig. 1e). The observed patterns of pairwise *F_ST_* suggest that isolation-by-distance likely drives differentiation between Mediterranean shearwater populations (Fig. 1e). These patterns were mirrored by the divergence statistic *d_XY_* (Fig. 1e). Interspecific divergence values were so low (*d_XY_* = 7.79 x 10^-4^ - 8.03 x 10^-4^) that they overlapped with levels of genetic diversity within populations (*π* = 7.58 x 10^-4^ - 7.83 x 10^-4^) (Fig. 1e). Indeed, the number of fixed SNPs between populations was remarkably low across all comparisons; for instance, only three SNPs out of ∼15M were fixed between MAL and MED (Supplementary Fig. 3). Overall, these results suggest that interspecific differentiation at the genomic level is shallow and is mostly driven by differences in allele frequencies.

### Divergence time estimates and demographic history of Mediterranean *Puffinus*

To accurately contextualise the events that shaped the evolution of Mediterranean *Puffinus,* we inferred their phylogenetic relationships and divergence times under the multispecies coalescent model (MSC)^21^. Coalescent trees and phylogenetic networks revealed very short internal branch lengths and a significant amount of reticulation, suggesting a recent and rapid diversification of all populations and the potential prevalence of gene flow (Fig. 2a; Supplementary Fig. 4). Divergence between Balearic and Yelkouan shearwaters was estimated at 19.9 ka (95% HPD: 12.6 - 30.3 ka), coinciding with the Last Glacial Maximum (LGM) (Fig. 2a; Supplementary Fig. 5a). Nevertheless, these estimates might underestimate the true divergence times because the applied model does not account for gene flow after divergence^22^. In contrast, this pattern was not replicated neither by mitochondrial data nor by the most differentiated genomic region between both taxa - hereafter scfF (see “Genome wide scans for signatures of divergent selection”). The tree topology of these regions reflected a deep split between Balearic and Yelkouan haplogroups dating back to the Middle Pleistocene: 551 ka (95% HPD: 349 - 765 ka) for mtDNA and 685 ka (95% HPD: 476 - 955 ka) for scfF (Fig. 2a; Supplementary Fig. 5b,c). We used forward simulations in SLiM^23^ to explore whether the topology and distribution of divergent haplotypes exclusive to low or no-recombining regions (e.g., mtDNA) could be explained by ancestral population structure and subsequent gene flow, as has been suggested elsewhere^24–26^. Indeed, coalescent trees resulting from simulations under the species’ modelled demographic history showed a similar topology as the trees inferred from empirical data: a middle Pleistocene divergence between mitochondrial haplogroups followed by recent diversification and migration events (Supplementary Fig. 6).

**Fig. 2:**
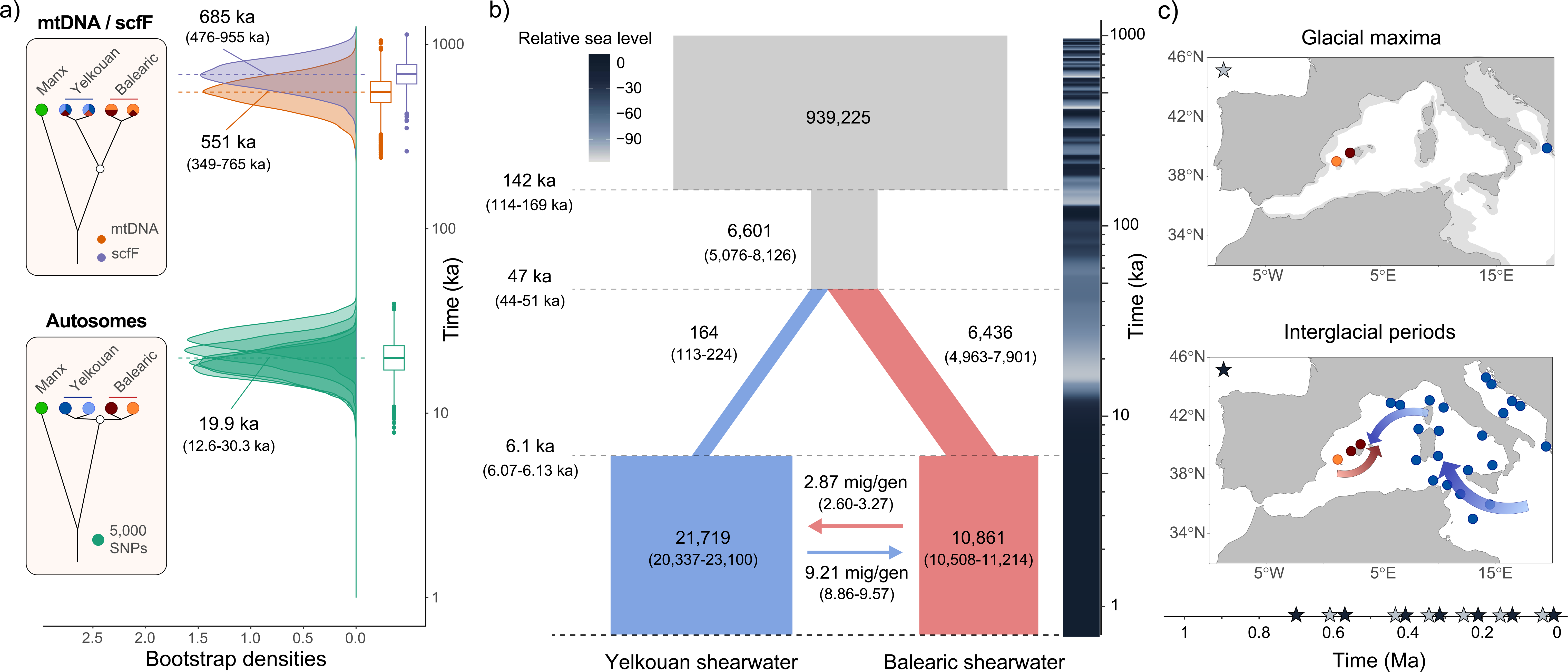
Glacial eustatic oscillations drive the evolutionary and demographic history of Mediterranean *Puffinus*. **a,** Divergence times estimated between major haplotypes/taxa inferred under the multispecies coalescent model using: i) Five independent sets of 5,000 genome-wide SNPs; ii) whole mitogenomes; and iii) 603 SNPs found within the centre of the selective sweep in scfF (see *Genome-wide scans for signatures of divergent selection*). Insets summarise the topologies inferred for each of the datasets and pie charts at the terminal branches are coloured based on the taxa as in Fig. 1. White dots indicate the node of interest whose divergence times are shown on the right. **b,** Joint demographic history of Balearic and Yelkouan shearwaters inferred using the unfolded 2D-SFS in ∂a∂i. Time is indicated in thousands of years (ka), population size in number of individuals and migration rates in number of migrants per generation. Numbers in parentheses show 95% confidence intervals. The colour-coded column on the right indicates the global average eustatic sea-level relative to the present-day level^101^. **c,** Schematic illustration of the hypothetical distribution and connectivity between *Puffinus* lineages in the Mediterranean during glacial and interglacial periods. Modern coastline is shown in dark grey; the coastline during the LGM is shown in light grey. Grey and black stars on the timeline represent glacial maxima and interglacial periods, respectively.

Given that hybridisation between Mediterranean *Puffinus* might affect our inference of divergence times under the MSC, we complemented the previous phylogenetic analyses by reconstructing the recent demographic history of these taxa using the two-dimensional site-frequency-spectrum (2D-SFS) between Balearic (PIT, MAL and CAB) and Yelkouan (MEN and MED) populations in ∂a∂i^27^. Our best fitting model supported a split between Balearic and Yelkouan shearwaters during the LGM, roughly 47.2 ka (95% CI: 43.5 - 51 ka) (Fig. 2b). After the split, both populations would have drifted in isolation until the Holocene interglacial, ∼6.1 ka (95% CI: 6.07 - 6.13 ka), when the model infers the onset of extensive gene flow between Mediterranean *Puffinus* (*m* = 2.97 - 9.21 migrants/generation) linked to a population expansion in both taxa (Fig. 2b). Population size oscillations linked to glacial cycles appear to have played a significant role in the demographic history of Mediterranean *Puffinus*, with larger population sizes inferred during interglacial periods than during glacial maxima (Fig. 2b,c). Indeed, the best model significantly outperformed models that did not account for census size changes both before and after the divergence of Balearic and Yelkouan shearwaters (Supplementary Fig. 7). The strikingly large ancestral population size inferred by the model is likely to be influenced by the existence of ancestral population structure^28^ during earlier glacial maxima. Periods of isolation, such as those that may have driven divergence in mtDNA and scfF, would result in genomic signatures akin to those expected under population expansions^29^.

### Characterisation of gene flow across Mediterranean *Puffinus*

We investigated whether the intermediate ancestries observed in Cabrera (CAB) and Menorca (MEN) resulted from introgression, rather than incomplete lineage sorting (ILS), by estimating *ƒ_4_* statistics across all possible population trios and performing *f*-branch tests (*ƒ_b_*)^30^. These tests supported introgression events between MEN and Balearic shearwaters, and between Yelkouan shearwaters and both CAB and MAL, albeit to varying degrees (Fig. 3a). In brief, the largest *ƒ_b_* values were observed between the geographically closest colonies, suggesting that the strength of gene flow correlates with geographic proximity. To explore whether gene flow also extends to the most distant populations (PIT and MED), we performed topology weighting across the genome using windows of 100 non-overlapping SNPs. For each window, we calculated the relative weight of each of the three possible topologies that include the main populations: PIT, MAL and MED (Fig. 3b). This analysis revealed that genome-wide proportions of these topologies were nearly identical: while the weight of the background topology - (MED,(MAL,PIT)) - was significantly higher than those of the alternative ones (FDR *q* < 1×10^-8^), there was no significant bias between the two alternative topologies (FDR *q* = 0.754; Fig. 3b). The absence of this bias contradicts the expectation that geographical distance should limit hybridisation and suggests that gene flow between the most isolated populations, PIT and MED, reaches similar magnitudes as between MAL and MED. The near absence of segregating SNPs between these three populations provides further support for this hypothesis (Supplementary Fig. 2).

**Fig. 3:**
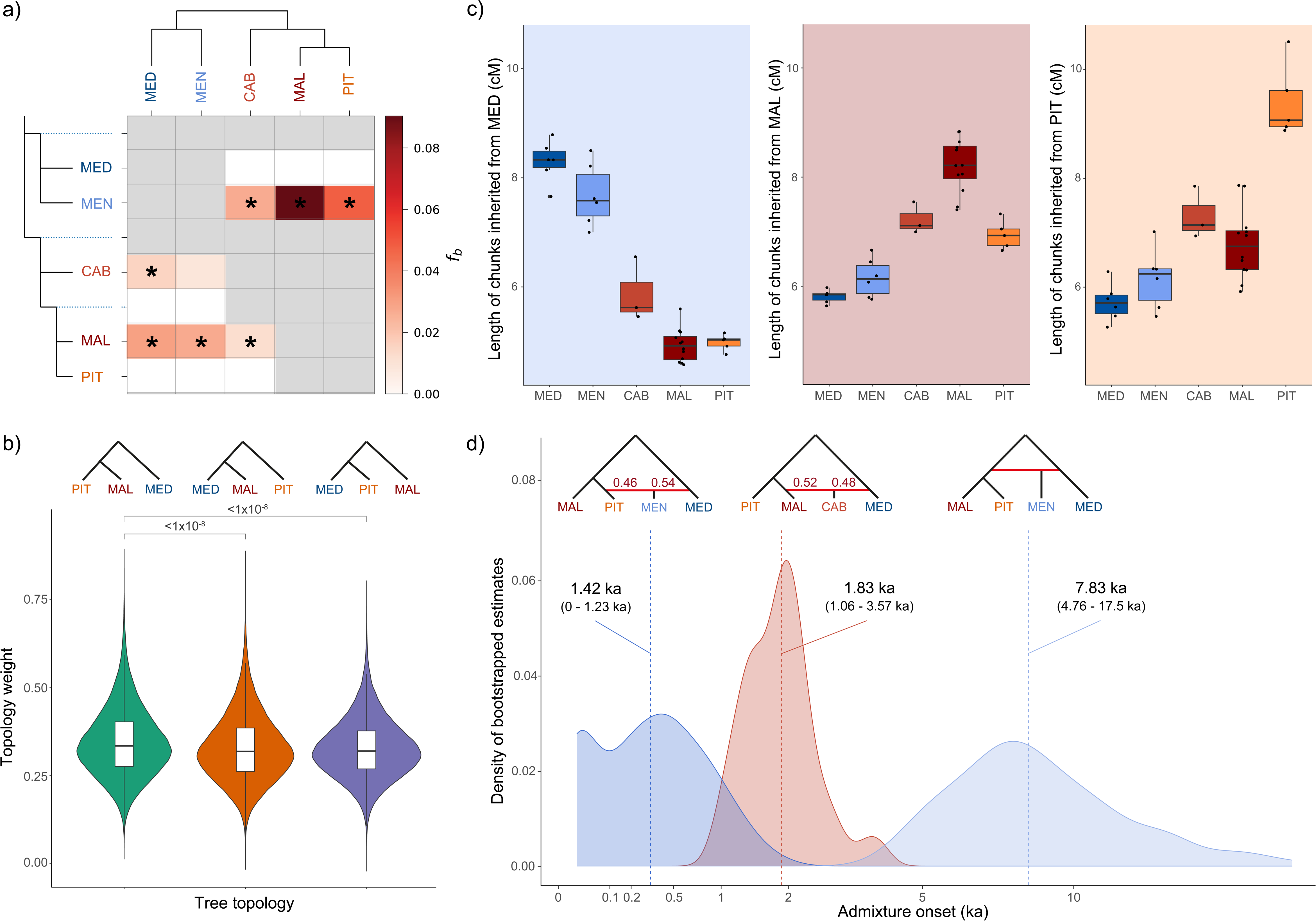
Widespread footprints of recent introgression between Balearic and Yelkouan shearwaters. **a,** Colour-coded heatmap according to the strength of introgression summarising the *f*-branch statistics estimated in Dsuite^30^. Dotted branches represent ancestral lineages within the phylogeny. Significant instances of excess allele-sharing are highlighted with an asterisk. **b,** Topology weights of the three possible topologies including only the three main *Puffinus* populations (PIT, MAL and MED) across 121,145 100-SNP windows. FDR-corrected p-values are shown for significant pairwise comparisons (Mann-Whitney *U*-test) **c,** Boxplots of ChromoPainter^89^ haplotype chunk lengths shared by target individuals (dots) and individuals from a donor population: MED (left), MAL (centre) or PIT (right). Boxplots show the median (centre), the first and third quartiles (box bounds), the smallest and largest values within 1.5x the interquartile range from the first and third quartiles (whiskers), and outliers (points). **d,** Density plots of estimates of admixture dates in thousands of years (ka) for the two populations showing higher levels of admixture (MEN in blue and CAB in red), using fastGlobetrotter^91^. Vertical dashed lines indicate the fastGlobetrotter onset estimates, while numbers in parentheses show 95% confidence intervals. Source populations and admixture proportions, when they could be inferred, are indicated in the schemes above each admixture event.

Additionally, we used phased haplotype information resulting from the combination of read-based and statistical phasing to characterise the sources and timing of admixture in MEN and CAB. First, we assigned each haplotype chunk across the genomes of individuals from these colonies to a parental population (PIT, MAL or MED). Individuals from MEN and CAB were inferred to inherit intermediate haplotype chunks from all parental populations (Fig. 3c). Moreover, the lengths of these chunks were shown to vary among hybrid individuals, as expected under recent introgression (Fig. 3c; Supplementary Table 3). Subsequently, we used these local ancestry paintings to infer the timing of admixture events that gave rise to MEN and CAB. The best-fit models indicated that ancestry patterns in MEN are best explained by multiple waves of admixture during the Holocene: a potentially ongoing wave of admixture involving PIT and MED (0.42 ka; 95% CI: 0 - 1.23 ka), and an older one between individuals of unknown source populations dating to 7.83 ka (95% CI: 4.76 - 17.54 ka) (Fig. 3d; Supplementary Table 4). Meanwhile, ancestry patterns in CAB are best explained by a single admixture event between MAL and MED during historical times (1.83 ka; 95% CI: 1.06 - 3.57 ka) (Fig. 3d; Supplementary Table 4). These findings support the hypothesis that population expansions after the LGM led to the establishment of hybrid colonies in secondary contact zones between Mediterranean *Puffinus*.

### Genome-wide scans for signatures of divergent selection

The combination of genome-wide scans for *F_ST_* and cross-population extended haplotype homozygosity (XP-EHH) statistics between Balearic and Yelkouan shearwaters allowed us to identify several shared outlier regions potentially involved in the phenotypic differentiation observed between these taxa (Fig. 4a; Supplementary Table 5). To understand the evolutionary mechanisms driving differentiation between shearwater taxa, we focused our efforts on the characterisation of scfF, the most differentiated *F_ST_* outlier region (Fig. 4). This genomic region is syntenic with macrochromosomal ends in related species (*Spheniscus humboldti* and *Ciconia maguari*), and exhibits long and distinct haplotypes that are unevenly distributed among *Puffinus* populations (Fig. 4b; Supplementary Tables 1,5). In particular, Yelkouan shearwaters are predominantly homozygous for haplotypes with alternate variants (H1), while Balearic shearwaters host both haplotypes at intermediate frequencies (Fig. 4b). The use of local PCAs and estimates of linkage disequilibrium (LD) revealed a gradual increase in recombination and a matching decline in population structure with increasing distance from the chromosome end, consistent with the expected footprints of a selective sweep in at least one taxon^31,32^ (Fig. 4c; Supplementary Fig. 8). We pinpointed the centre of the selective sweep within a 200 kb region with homogeneous LD and local population structure patterns, which encompasses several genes (Fig. 4c,d; Supplementary Fig. 8). Among these genes, several have been identified in previous studies as responsible for phenotypic changes in plumage colouration (e.g., *AP1G1*^33^), social behaviour (e.g., *ATXN1L*^34^) or pre-migratory hyperphagia (e.g., *TAT*^35,36^) in other species (Supplementary Table 6). We found four non-synonymous substitutions between H0 and H1 haplotypes. Notably, one of these substitutions was identified in a tyrosine aminotransferase (*TAT*) whose coding region is highly constrained across a 364-way bird genome alignment (Fig. 4d; Supplementary Fig. 9).

**Fig. 4:**
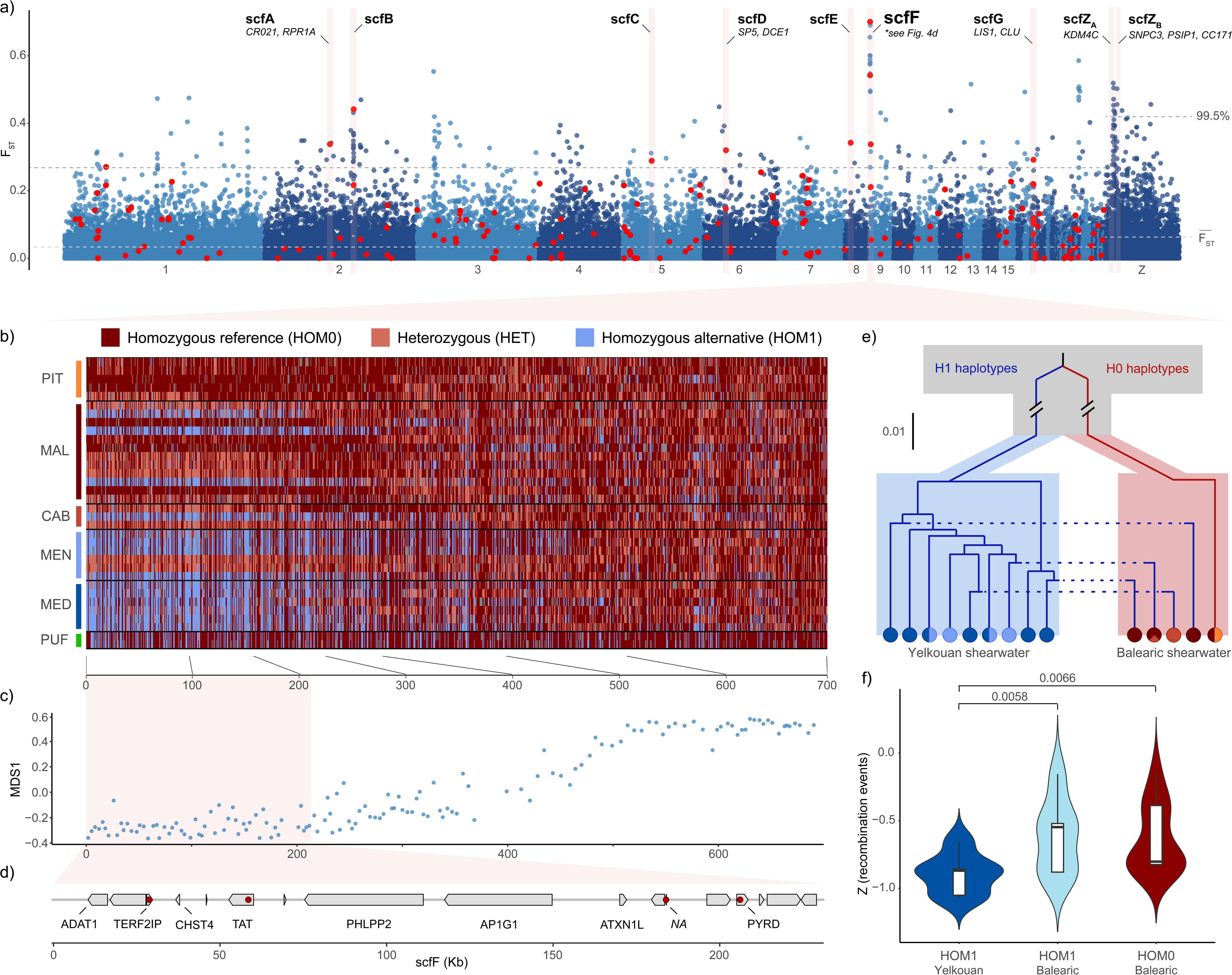
Gene flow has led to the introduction of Yelkouan adaptive haplotypes into Balearic shearwaters. **a,** Genome-wide scans for pairwise *F_ST_* between Balearic and Yelkouan shearwaters using 25 kb non-overlapping sliding windows. Red dots highlight windows with outlier values (> 99.5% of all windows) for XP-EHH. Autosomal windows with shared outliers for both statistics and sex-linked scaffolds with consecutive *F_ST_* outliers are highlighted as candidate regions driving interspecific differentiation. Genes within candidate windows are also shown. Dotted lines indicate the mean and 99.5% outlier threshold across autosomes and the Z chromosome. Scaffolds are ordered according to synteny with the Humboldt penguin reference genome. **b,** Genotype plot of a representative candidate window - scfF - polarised for the major allele in PIT. The genotype of each SNP is depicted as heterozygous (HET: light red), homozygous reference (HOM0: dark red) or alternative (HOM1: light blue). **c,** Multidimensional scaling (MDS) of distances between local PCA maps (dots) using 25 SNP windows across scfF; distances using the MDS1 axis are shown. **d,** Annotated genes across the centre of the selective sweep within scfF, with selected genes indicated. Red dots illustrate non-synonymous SNPs that segregate between haplotypes. **e,** Neighbour-joining tree of whole phased scfF haplotypes of homozygous individuals visualised over a schematic representation of the demographic history of Mediterranean *Puffinus*. Pie charts are colour-coded according to Fig. 1 and indicate the proportion of individuals of each population in any given clade. **f,** Recombination rates of H0 and H1 scfF haplotypes in homozygous Balearic and Yelkouan shearwaters, measured as the normalised results of 4-gamete tests across 25 kb windows. FDR-corrected p-values are shown for significant pairwise comparisons (Mann-Whitney *U*-test).

Next, we investigated the origin of the intermediate haplotype frequencies in scfF in Balearic shearwaters. Balancing selection acting on ancestral haplotypes can preserve genomic modules at intermediate frequencies, enabling rapid adaptation to changing selective pressures in a given population^37^. Given that the divergence between H0 and H1 haplotypes predates speciation (Fig. 2), we explored the balancing selection hypothesis by analysing Tajima’s *D* and allele frequency correlation (*β*) along scfF. These summary statistics showed significantly higher values in scfF compared to the genome-wide distribution (Mann-Whitney *U*-test, *p_Tajima’s D_* = 0.033; *p_ß_* = 2.46*10^-8^), an expected outcome under balancing selection^38^ (Supplementary Fig. 10). However, when we investigated tree topologies between phased scfF haplotypes, we observed multiple coalescent events between H1 haplotypes from Balearic and Yelkouan shearwaters (Fig. 4e; Supplementary Fig. 11). Such a pattern is unlikely to arise from ancestral balancing selection, which should result in the monophyly of H1 haplotypes, and it is better explained by multiple recent introductions of H1 haplotypes from Yelkouan into Balearic shearwaters. The comparison of LD estimates between homozygous reference (HOM0) and alternate (HOM1) individuals of both taxa, measured using standardised *ZnS* and 4-gamete test values, revealed significantly lower recombination rates within scfF in HOM1 Yelkouan shearwaters compared to both HOM1 (Mann-Whitney *U*-test, FDR *q_4gam_* = 0.006) and HOM0 Balearic shearwaters (Mann-Whitney *U*-test, FDR *q_4gam_* = 0.007) (Fig. 4f; Supplementary Fig. 12 & 13). Alongside this, we highlight two further sources of evidence that suggest HOM1 genotypes are only positively selected in Yelkouan shearwaters: the latter species shows lower genetic diversity (*π*) and lower Tajima’s *D* values when compared to Balearic shearwaters within scfF, albeit comparisons were not significant (Supplementary Fig. 12). Our findings suggest that, although H1 haplotypes are not necessarily adaptive in Balearic shearwaters, they have been recurrently introduced through hybridisation from Yelkouan shearwaters. Overall, this highlights that under pervasive introgression, even adaptive alleles can be exchanged between populations.

### Estimates of genetic diversity, inbreeding and genetic load

The contemporary small population size and recent demographic collapse in Balearic shearwaters are expected to result in pronounced genomic footprints of inbreeding, which could further imperil the species’ survival. In contrast to these demographic observations, we detected similar levels of genome-wide heterozygosity across Balearic and Yelkouan shearwater individuals (*H_O_ =* 1.70×10^-3^ - 1.97×10^-3^), with no significant differences between populations (Fig. 5a; Supplementary Table 7). A similarly uniform pattern was found for inbreeding coefficients, measured as the proportion of the genome in runs of homozygosity (*F_ROH_*) over 100 kb in length (Fig. 5b; Supplementary Table 7). We only detected significant differences in pairwise comparisons between Balearic populations: long ROH (ROH >1 Mb; *F_LROH_*), which are indicative of more recent inbreeding events, were significantly more abundant in MAL (*F̅_LROH,1Mb_* = 1.03×10^-2^) than in PIT (*F̅_LROH,1Mb_* = 0.45×10^-2^; Mann-Whitney *U*-test, FDR *q* = 0.037; Fig. 5b). Both individuals of Manx shearwater (used as the outgroup) showed slightly higher heterozygosity than Mediterranean *Puffinus* (*H_O_ =* 2.01×10^-3^ - 2.48×10^-3^), yet inbreeding coefficients were also higher (*F̅_ROH,100kb_* = 8.03×10^-2^; *F̅_ROH,1Mb_* = 2.49×10^-2^). This suggests that Mediterranean *Puffinus* have not suffered severe population bottlenecks compared to congeneric shearwater taxa with larger populations (Supplementary Table 7).

**Fig. 5:**
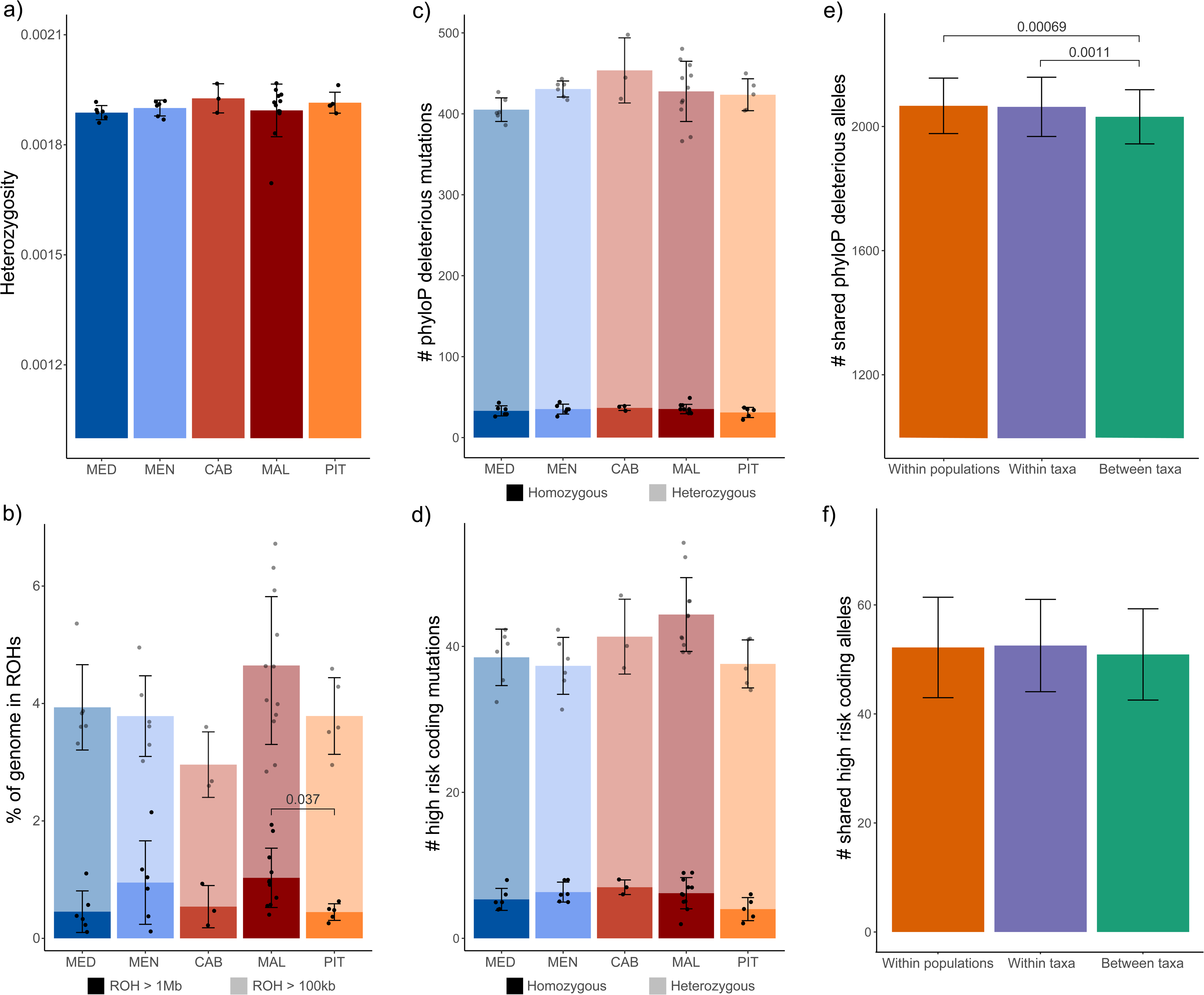
Reduced footprints of inbreeding and shared genetic load across Mediterranean *Puffinus* populations. **a,** Average genome-wide heterozygosity per individual. **b,** Individual inbreeding coefficients based on ROH (*F_ROH_*). Dark bars show the proportion of the genome in ROH ≥ 1 Mb; light bars show the remaining proportion found in ROH ≥ 100 kb. **c,d,** Proxies of genetic load: number of evolutionary constrained mutations inferred with phyloP^39^ **(c)** and number of high-risk coding variants per individual, as estimated by snpEff^40^ **(d)**. Dark bars show the number of variants found in homozygosity; light bars show those found in heterozygosity. **e,f,** Estimates of shared genetic load between pairs of individuals found in the same population, in the same taxon or in a different taxon: number of shared deleterious alleles between individuals according to phyloP **(e)** and snpEff **(f)**. In all cases, error bars indicate standard errors and FDR-corrected p-values are shown for significant pairwise comparisons (Mann-Whitney *U*-test or Welch’s t-test - see *Methods*).

To further investigate the potential effects of inbreeding in Balearic shearwaters, or lack thereof, we quantified genetic load using proxies based on evolutionary conservation across the avian phylogeny (phyloP^39^) and functional effect prediction (snpEff^40^). Similar to our comparisons of genetic diversity, both approaches failed to recover significant differences in the amount of genetic load between any Mediterranean *Puffinus* populations (Fig. 5c,d; Supplementary Table 7). The SFS of both (i) SNPs with highly-conserved phyloP scores and (ii) SNPs with high risk snpEff annotations were skewed towards low-frequency variants compared to neutral positions (Supplementary Fig. 14). Such patterns are expected if the annotated SNPs cause a reduction in fitness^8^. Overall, we argue that the striking lack of differentiation in genetic diversity and genetic load among populations with drastically different population sizes can be explained by the exchange of divergent genetic material through continuous and pervasive hybridisation. This hypothesis is supported by the absence of significant differences in the amount of coding deleterious mutations annotated by snpEff shared within each population compared to those shared between populations from different taxa (t-test, FDR *q* = 0.079; Fig. 5f). Nevertheless, this pattern is not shared by phyloP-scored highly-conserved SNPs, as conspecific populations share more deleterious mutations than allospecific ones (t-test, FDR *q* = 1.05×10^-3^; Fig. 5e).

### Forward simulations to assess the future risk of inbreeding

Our findings suggest hybridisation is crucial to mitigating inbreeding depression in Balearic shearwaters. Thus, we set out to explore the effects of introgression on genetic diversity and population size under realistic future demographic scenarios by employing forward simulations in SLiM^23^ using 25 Mb simulated diploid chromosomes. We compared the outcome of three possible migration scenarios: a) no hybridisation due to the local extinctions of MEN and CAB; b) current migration rates (*m = 10 migrants/generation*); and c) potential human-mediated translocation from Yelkouan colonies (*m = 100 migrants/generation*). Moreover, we contrasted the results under three demographic scenarios using life-history parameter estimates outlined in a recent viability analysis^12^: i) current demographic projections; ii) demographic projections assuming increased survival probabilities due to immediate bycatch reduction; iii) projections if the implementation of bycatch reduction measures is delayed by 25 years (Supplementary Tables 8 & 9). Balearic and Yelkouan shearwater populations were simulated using current estimates of population size, population differentiation (*F_ST_*) and heterozygosity.

Our simulated projected population sizes matched predictions from previous viability analyses^12^: under the current demographic projections and a lack of migration, we inferred a mean time to extinction of 59.7 years (95% CI: 49.9 - 74.2 years) for the Balearic shearwater (Fig. 6a). Under this model, increased migration rates only delayed the projected extinction times by a maximum of two decades (Supplementary Fig. 15). On the other hand, bycatch reduction resulted in the survival of Balearic shearwaters past the 500-year mark, albeit with varying population sizes. We found that the “immediate bycatch reduction” scenario resulted in hundreds of surviving individuals after 500 years (*n̅* = 575 - 774 depending on migration rates; Fig. 6a,b). Conversely, a delay in such measures would result in markedly reduced population sizes at that time, especially in the absence of migration (*n̅* = 3; 95% CI: 0 - 12; Fig. 6a,b).

**Fig. 6:**
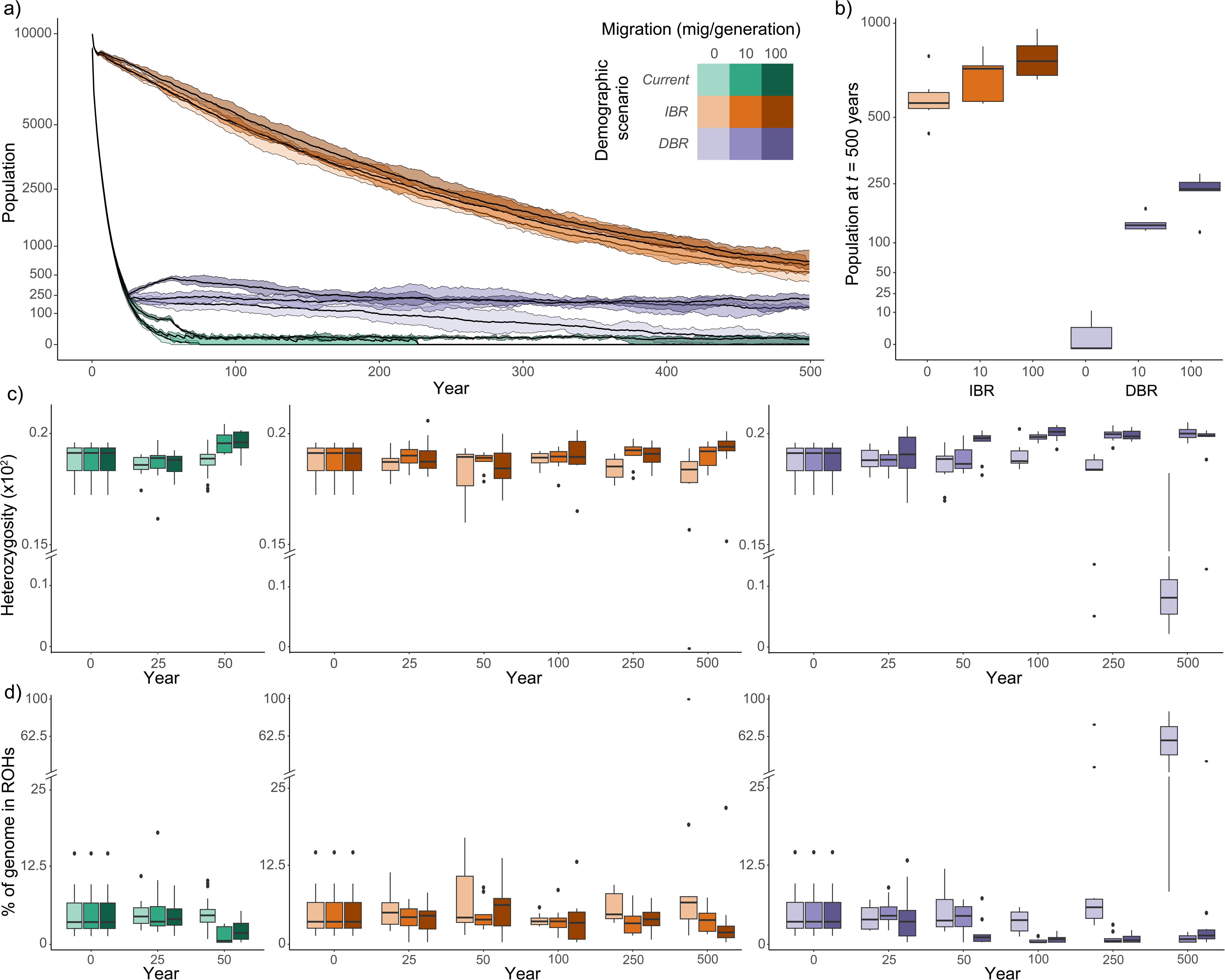
Forward simulations of population size and genetic diversity under future demographic projections in Balearic shearwaters. **a,** Projection of simulated census sizes for Balearic shearwaters under nine potential future demographic scenarios. Scenarios illustrate variation in migration rates between Balearic and Yelkouan shearwaters (*m* = 0, *m* = 10 and *m* = 100 migrants/generation) and in demographic parameters related to bycatch mitigation strategies under: i) current scenario; ii) immediate bycatch reduction (IBR); and iii) delayed bycatch reduction (DBR). Black lines represent the average projected census size across five SLiM^23^ simulations; shaded ribbons indicate 95% confidence intervals. Legend shows the colour codes for each combination of demographic scenario and migration rate, with migration rates indicated in migrants per generation. **b,** Projected census size of the Balearic shearwater after 500 years under different combinations of demographic scenarios and migration rates. **c,** Average simulated genome-wide heterozygosity per individual at multiple time points under nine different combinations of demographic scenarios and migration rates. **d,** Simulated individual inbreeding coefficients based on ROH over 100 kb in length (*F_ROH_*) across demographic scenarios. In **(b)**, **(c)** and **(d)**, different colours indicate different demographic scenarios, while increasing opacity reflects increasing migration rates as shown in the legend in **(a)**). In all cases, boxplots show the median (centre), the first and third quartiles (box bounds), the smallest and largest values within 1.5x the interquartile range from the first and third quartiles (whiskers), and outliers (points).

Indeed, the only significant drop in heterozygosity compared to starting values (Mann-Whitney *U*-test, FDR *q* = 2.84×10^-4^) was observed after 500 years in the “delayed bycatch reduction” scenario under no migration (Fig. 6c). This was also associated with a rise in *F_ROH_* (Mann-Whitney *U*-test, FDR *q* = 1.62×10^-4^; Fig. 6d). In all other cases, from 100 years onwards increasing migration also resulted in higher values of heterozygosity and a decrease in *F_ROH_*, although differences were not significant (Fig. 6c,d). Overall, these simulations suggest that inbreeding depression does not constitute an immediate problem for Balearic shearwaters. Instead, inbreeding would only become a threat to the species’ long-term survival under a combination of: a) a delayed implementation of bycatch-reduction measures and b) the loss of migratory connectivity with Yelkouan shearwaters (i.e. the extinction of Menorcan and Cabreran populations).

## DISCUSSION

Our research provides a key case study to examine the evolutionary significance of hybridisation and its conservation implications in a Critically Endangered flagship species. We present a comprehensive overview of the evolutionary and demographic processes that have shaped the genomes of Mediterranean *Puffinus* species. Our results uncover very low levels of genome-wide differentiation between Balearic and Yelkouan shearwaters, a phenomenon that we attribute to the extensiveness of gene flow between them throughout their evolutionary history. While hybridisation appears to be most prevalent in the islands of Menorca and Cabrera, as previously suggested^13,14,41^, its effects permeate even to the most differentiated populations (Fig. 3). Migration rates inferred through demographic modelling greatly exceed values typically assumed to ensure intraspecific population connectivity^42,43^. Indeed, differentiation between these taxa seems primarily driven by differences in allele frequencies associated with local adaptation, and even these alleles can be exchanged between taxa. This shows that interspecific gene flow has been a major component in the evolutionary history of these taxa and should be taken into account in the ongoing debate about the specific status of the Balearic shearwater^18^.

The complementary use of time-calibrated phylogenies and demographic modelling has enabled us to support the hypothesis^19,44^ that eustatic oscillations during Pleistocene glacial cycles were the main drivers of demographic changes and population divergence in Mediterranean *Puffinus*. The drop in sea-level during glacial maxima caused the Sicilian channel to become narrower, shallower, and highly saline^45–47^. Consequently, the habitat became unsuitable for foraging shearwaters, leading to allopatric differentiation on either side of the channel. During interglacial periods, population expansions driven by the loss of geographic barriers and increased habitat availability likely facilitated the widespread hybridisation between shearwater taxa through secondary contact, leading to speciation reversal, as exemplified in other taxa^48^. The repetition of this process through glacial cycles has likely resulted in highly-differentiated haplotypes with deep coalescence times being maintained only in low to no-recombining regions, such as mtDNA and scfF (Fig. 2). Under current climatic projections, widespread gene flow between Mediterranean *Puffinus* species is likely to persist. This case draws attention to the role of climate change in driving macroevolutionary processes such as speciation reversal.

The pervasiveness of gene-flow between Balearic and Yelkouan shearwaters has significantly eroded the genomic landscape of differentiation, with the notable exceptions of a few outlier loci, best exemplified by the scfF (Fig. 4). Within this region, divergent haplotypes were likely maintained by a selective sweep in at least one taxon, most likely in the Yelkouan shearwater, as evidenced by its reduced values of recombination rate, genetic diversity and Tajima’s D. Remarkably, the same genomic region has been identified as a potential driver of differentiation between resident and migratory phenotypes in songbird species^49^. Within this region, *TAT* emerges as a promising candidate gene for driving ecological and/or phenotypic differences between Balearic and Yelkouan shearwaters linked to their distinct migratory strategies. This gene is known to play a significant role in the liver and brain of songbirds during the fattening stage that precedes migration^35,36^. The ancient coalescent times between scfF haplotypes suggest that different migratory strategies might have originated in response to Pleistocene climatic shifts. The current exchange of alleles under selection due to widespread hybridisation, as evidenced in this study, is expected to enhance the adaptive potential of Mediterranean *Puffinus* to climate change through changes in their migratory behaviour.

Our study adds onto the growing body of research emphasising the beneficial role of hybridisation in maintaining genetic diversity and avoiding inbreeding depression^3–5,50,51^. Shearwaters offer a compelling example of how introgression needs to be carefully considered as a potential source for genetic rescue in depleted populations, especially in cases where hybridisation has been widespread throughout a species’ evolutionary history. Through forward simulations we uncover that the preservation of genetic connectivity between Balearic and Yelkouan shearwaters would mitigate the effects of inbreeding depression (i.e. natural genetic rescue), especially in face of bottlenecks caused by a delay in the implementation of protection measures. If hybrid colonies go extinct (i.e. Menorca and Cabrera), translocation attempts could serve as an additional way to boost population size while guaranteeing the maintenance of genetic diversity in the Balearic shearwater. These actions have been successful in other shearwater taxa^52–54^. Nevertheless, we show that focusing only on actions to preserve genetic diversity is insufficient to ensure the survival of the Balearic shearwater. While the protection of hybrid populations and even translocation attempts can help to avoid the effects of inbreeding depression in the future, current demographic projections indicate that Balearic shearwaters will go extinct in the next 100 years regardless of these measures. Only via concerted conservation efforts, especially those aimed at rapidly reducing bycatch-driven mortality, can we prevent the imminent extinction of the Balearic shearwater.

## MATERIALS AND METHODS

### Sampling, library preparation and sequencing

Genomic DNA was extracted from blood samples of 36 individuals sampled across the distribution of Balearic and Yelkouan shearwaters, as well as two individuals of their outgroup: the Manx shearwater (Figure 1a; Supplementary Table 1). Samples were preserved in 100 % ethanol and kept at −20°C. We included 3-12 individuals from each of the sampled Balearic shearwater populations: Pitiüses, Cabrera, Mallorca and Menorca and single individuals from six colonies of Yelkouan shearwater (Figure 1a; Supplementary Table 1). DNA was extracted using DNeasy Blood & Tissue Kit (Qiagen) following the manufacturer’s instructions. Both Illumina TruSeq DNA PCR-Free and TruSeq Nano DNA Kit libraries (insert-size = 350 bp) were prepared by Macrogen (South Korea) and sequenced using two HiSeq X runs for paired-end reads (2x150bp) (Supplementary Table 2).

### Variant calling and filtering

Quality control was performed using FastQC v0.11.7 (https://www.bioinformatics.babraham.ac.uk/projects/fastqc/) and adapter sequences were removed with Trim Galore v0.4.5 (https://www.bioinformatics.babraham.ac.uk/projects/trim_galore/). Clean reads were mapped to the Balearic shearwater reference genome^19^ using bwa mem v0.7.7^55^ and duplicate reads were removed using MarkDuplicates in PicardTools v2.20.4 (http://broadinstitute.github.io/picard/). Variant calling was performed using both HaplotypeCaller in GATK4 v4.1.9^56^ and Freebayes v1.2^57^. We then created an intersect file from the GATK and Freebayes VCF files using bcftools v1.8^58^, keeping only those variants called by both GATK and Freebayes. We set the following hard filtering parameters to exclude low quality variants in GATK: QD < 2.0, FS > 60.0, MQ < 20.0, and ReadPosRankSum < – 8.0. Relatedness statistics between individuals were estimated using VCFtools v.0.1.15 (option --relatedness2)^59,60^. For any 2nd-degree or more closely related pair of individuals, one was excluded from the raw VCF file to avoid biases in downstream analyses^60,61^, resulting in the removal of two individuals (Supplementary Table 1). We also removed low-confidence variants by applying the following filters: only biallelic SNPs filtered for quality (--minQ 30) and coverage (--minDP 4, --maxDP 50) using VCFtools v.0.1.15^59^ were retained in the final VCF files. Depending on the needs of each analysis, separate VCF files were generated using different cutoffs for completeness, minimum allele count and LD-based pruning (Supplementary Table 10). Finally, we also masked sites found within the mappability and repeat masks of the Balearic shearwater reference genome (see Cuevas-Caballé *et al.* 2022^19^).

We repeated the variant calling process with reads that mapped exclusively to the reference mitochondrial genome of the Balearic shearwater, using the haploid variant calling mode of Freebayes v1.2^57^ and different quality and coverage cutoffs (--minQ 750; --min-meanDP 41, --max-meanDP 55, --minDP 9) based on the empirical distributions of these variables in mitochondrial SNPs. As sex chromosomes are known to potentially reflect different evolutionary histories than autosomes^62^, we also divided variants for autosomes and sex chromosomes into different files for downstream analyses. We determined the putative sex-linked scaffolds in the reference genome for the Balearic shearwater using a custom pipeline that relies on: a) reduced coverage in the resequenced females compared to males; and b) conserved synteny with sex chromosomes of the phylogenetically closest chromosome-level genome assembly available (Humboldt penguin (*Spheniscus humboldti*)) using pairwise minimap2 v2.11^63^ with default parameters.

### Population structure analyses

To visualise population structure, we performed principal component analyses (PCAs) in PLINK v1.90b5.3^64^ after pruning linked sites with an R^2^ > 0.1. We inferred clusters and individual ancestry proportions using the Struct-f4 Rcpp package^20^ for all populations between *K* = 1 and *K* = 5. By using f4 statistics instead of individual allele frequencies, Struct-f4 reduces biases caused by the amount of genetic drift exclusive to single populations, which affects clustering methods that assume Hardy-Weinberg equilibrium, such as ADMIXTURE and STRUCTURE^20,65^. To further explore the observed patterns, we calculated average genome-wide *F_ST_* values between all population pairs using VCFtools v.0.1.15^59^. Furthermore, we estimated the absolute divergence between populations (*d_xy_*) and within-population nucleotide diversity (*π*) in 25 kb non-overlapping sliding windows across the genome using the PopGenome v.2.7.5 R package^66^ and adjusted the denominators based on the proportion of masked base-pairs within each window. Fixed SNPs differing between population pairs were also obtained through the comparison of VCFs using custom scripts. All analyses were performed separately for the autosomal and sex-linked datasets.

### Estimating divergence times between Balearic and Yelkouan shearwaters

To visualise phylogenetic relationships across our samples, we first calculated pairwise distances under the Kimura two parameter substitution model and then inferred neighbour-joining phylogenetic trees using the *dist.dna* and *njs* functions implemented in the ape v.5.6-2 R package^67^. We also visualised reticulation across the phylogeny by constructing a splits network using the Neighbor Net agglomerative algorithm on SplitsTree5 v.5.0.0_alpha^68^ with the aforementioned pairwise distance matrix.

We inferred divergence times between *Puffinus* populations under the multispecies coalescent model using the SNAPP add-on package for BEAST2^21^ using the relationships inferred in the splits networks as the starting topologies. Due to the high computational demands of SNAPP, we first performed this analysis only with a reduced set of 5000 randomly selected genome-wide SNPs and 3 individuals per population. To account for the stochasticity in the SNP selection step, we repeated this procedure using ten different datasets of 5,000 random SNPs. We removed all singletons from the SNP-selection step to minimise the effect of drift within individuals. The input files for SNAPP were prepared with the script snapp_prep.rb^69^, which implements a strict-clock model and a pure-birth tree model, and the analyses were run for 500,000 MCMC iterations. The use of a strict-clock model is justified when using genome-wide data of closely related taxa because substitution rates are not expected to vary significantly between lineages^69^. We used a normal distribution to constrain the root age of the tree based on past estimates of the divergence time between the Manx shearwater and its Mediterranean congeners (mean = 2.084 Ma; standard deviation = 0.304)^18^.

Neighbour-joining phylogenies using mtDNA and scfF show remarkably different topologies from the genome-wide trees: instead of a recent diversification, individuals segregate into two distinct haplogroups. To date the divergence between haplogroups, we followed a similar approach to Matschiner *et al.* (2022)^70^ by repeating the SNAPP runs for 603 biallelic SNPs found within scfF using the same parameters as in the genome-wide runs. We included six homozygous individuals per haplogroup divided into two clades based on the topology recovered when including all individuals (Supplementary Fig. 4). We excluded singletons analogously to the genome-wide runs. In the case of mtDNA, we dated the divergence between haplogroups using whole mitogenomic sequences and conducted MCMC runs for 10,000,000 generations in BEAST 1.10.4^71^ implemented with BEAGLE^72^. We constrained the root age of the tree with the same parameters as in the SNAPP runs and applied a HKY substitution model and a yule speciation model. In all runs, convergence of parameter estimates was assessed through the visualisation of trace files in TRACER v1.7.2^73^ and the trees (excl. the first 15% of MCMC runs) were summarised with TREEANNOTATOR v1.10.4^74^.

### Demographic history inference and demographic modelling

To investigate the demographic history of Mediterranean *Puffinus*, we generated the folded SFS for both Balearic (PIT, MAL and CAB) and Yelkouan shearwaters (MEN and MED), as well as a folded 2D-SFS, using ∂a∂i^27^. Through the comparison of the 2D-SFS, we observed differences in datasets depending on the genomic features excluded (none *vs* genes *vs* exons) or the chromosomes included (all autosomes *vs* macrochromosomes) likely caused by deviations from the neutral demographic history (Supplementary Fig. 16). Consequently, in the SFS used in downstream demographic analyses we only included SNPs found in scaffolds syntenic with macrochromosomes in *S. humboldti* and excluded all positions found within genes.

We first attempted to infer the demographic history of each taxon using StairwayPlot2^75^ based on the folded SFS of three datasets: a) Balearic shearwaters; b) Yelkouan shearwaters; and c) all Mediterranean *Puffinus.* However, sampling within a deme that has suffered extensive gene-flow from other demes is known to heavily bias demographic inference, both in the estimates of timing and population size^29^. Indeed, results from demographic inference using StairwayPlot were found to be unreliable, presumably due to the effects of gene-flow: demographic histories with each of the three datasets were nearly identical, suggesting a high similarity between the SFS of all populations (Supplementary Fig. 17).

Consequently, we performed further demographic analyses and model testing using the 2D-SFS with ∂a∂i v2.3.2^27^ as implemented in dadi-cli v0.9.2^76^ to model the effects of potential structure and gene flow. We used a hierarchical approach for model choice, first comparing simple models and then building on the best simple model by testing whether the addition of certain parameters led to a significantly better likelihood. We first compared the six demographic models that represented the plausible recent divergence scenarios between Balearic and Yelkouan shearwaters, ignoring population size changes: a) isolation without migration, b) isolation with continuous symmetric or c) asymmetric migration; d) secondary contact with symmetric or e) asymmetric migration; and f) split with only initial migration (Supplementary Fig. 6a). We built on the best model by comparing the effects of population size changes during different points of the demographic history: a) before the split, b) after the split, and c) both before and after the split (Supplementary Fig. 6b). To maximise the robustness of the inferences^77^, we repeated all models with the unfolded 2D-SFS (Supplementary Fig. 6). We polarised the unfolded SFS using the inferred ancestral states between Balearic and Cory’s shearwaters (*Calonectris borealis*) in a 364-way bird genome alignment (see “Estimates of genetic load using evolutionary conservation”) using the *halBranchMutations* function of the HAL toolkit^78^.

We optimised the parameters of each model by conducting a minimum of ten parallel runs, with 100 optimisation rounds each in dadi-cli and subsequently checking for convergence using the *dadi-cli BestFit* option. If convergence was not reached, we used the optimised parameter estimates as starting values for an additional five runs, a process that was repeated until convergence was achieved. In parallel, we generated 100 block-bootstrapped SFS by sampling 10 Mb genome chunks with replacement. Pseudo-replicates were then used to estimate parameter uncertainty for the best model with Godambe Information Matrix to estimate 95% confidence intervals using a step size of 0.001 as described in the ∂a∂i guidelines^27^. Furthermore, to avoid potential biases in model likelihoods caused by unaccounted linkage, model choice between nested models was based on the significance of likelihood-ratio tests (LRT) between the optimal model and the next best one. We assessed the significance of the LRT while controlling for Type I error by normalising the difference in log-likelihoods as outlined in Coffman *et al.* (2016)^79^ (Supplementary Fig. 7).

In all demographic analyses we scaled the inferred parameters using a mutation rate of *μ* =1.27×10^-8^ mutations per nucleotide per generation; total sequence length of *L* = 4.86×10^7^ bp; and a generation time of *g* = 12.8 years^12^. The total sequence length was calculated as:

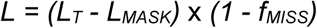

where *L_T_* is the total genome length, *L_MASK_* is the overlapping length of regions masked when generating the SFS (genes; scaffolds syntenic with microchromosomes and sex chromosomes; and regions masked during the SNP filtering process - see “Variant calling and filtering”) and *f_MISS_* is the proportion of SNPs with missing data that are lost when generating the 2D-SFS. As body size can have an effect on mutation rate, we fitted a logarithmic regression between body mass and mutation rate to the core waterbird species for which a germline mutation rate has been quantified^80^ (Supplementary Fig. 18). We then intercepted the average weight of Balearic shearwater (500g) to predict the mutation rate used in these demographic analyses.

### Forward simulations to assess the origin of mito-nuclear discordance

We assessed the feasibility that deep mitochondrial haplogroups originate from ancestral population structure by performing individual-based forward simulations recapitulating the demographic history of Mediterranean *Puffinus* in SLiM v4.0.1^23^. Using the Wright-Fisher (WF) implementation of SLiM, we simulated full mitogenomes reconstructing the last 800,000 years of Mediterranean *Puffinus* evolutionary history (Supplementary Fig. 6a). For these simulations, we modelled only the female individuals of the two populations and ensured that the mitochondrial genome, consisting of a 16,000 bp non-recombinant DNA fragment, was clonally inherited from one generation to the next. Coalescent tree sequences were output at the end of each simulation (cycle 800,000). These included single trees due to the lack of recombination in mtDNA. Mutations were generated along the coalescent tree with msprime v.1.2.0^81^ using a mutation rate of 1×10^-8^ mut/site/year. Finally, we output a VCF including 20 individuals of each taxon using tskit v.0.5.6

We first run a burn-in step of 235,000 cycles to obtain a fully coalesced population, *p_0_*, by simulating a panmictic *Puffinus* population with a carrying capacity of *K_0_* = 35,000 individuals. To simulate ancestral population structure, at cycle 235,000 we simulated a split of *p_0_* into two equally-sized subpopulations representing Balearic and Yelkouan mtDNA lineages, *p_1_* and *p_2_* respectively, by randomly drawing individuals from *p_0_* and moving them into one of the two subpopulations. We set the carrying capacity of each subpopulation to the current *N_e_* estimates obtained through demographic modelling: *K_1_* = 7,000 individuals for *p_1_* and *K_2_* = 14,000 individuals for *p_2_* (Supplementary Fig. 6a). We simulated no exchange of migrants or population size changes for the next 435,000 cycles. During the last 130,000 cycles we simulated the effects of glacial cycles on population size and migration, as previously inferred with demographic modelling - namely, population expansions and increased migration in interglacial periods, and bottlenecks in isolation during glacial maxima. To do that, we first simulated the effects of the Last Interglacial: at cycle 670,000 we simulated a demographic expansion in both subpopulations (*K_1_* = 35,000 individuals; *K_2_* = 70,000 individuals) coupled with the onset of migration using estimates of current migration rates obtained with ∂a∂i (9 migrants/generation from *p_2_* to *p_1_*; 2 migrants/generation from *p_1_* to *p_2_*). Migrants were randomly drawn from a uniform distribution of age classes (3-49 years) to include only adult individuals. Next, we simulated the effects of the Last Glacial Period by simulating a bottleneck in both populations (*K_1_* = 4,200 individuals; *K_2_* = 140 individuals) with no gene flow between cycles 685,000 and 794,000. Finally, after cycle 794,000 we simulated a last population expansion and secondary contact during recent times using the ∂a∂i estimates of current population sizes and migration rates (Supplementary Fig. 6a).

To understand whether divergence ages between haplogroups of simulated sequences were consistent with empirical data, we built time-calibrated phylogenies in BEAST 1.10.4^71^ implemented with BEAGLE^72^. We used the substitution rate inferred in the empirical runs with *Puffinus* mitogenomes as a prior by using a normal distribution centred on 3.82×10^-3^ substitutions/bp/Myr (standard deviation = 8×10^-4^). We applied a HKY substitution model, a yule speciation model and conducted MCMC runs for 10,000,000 generations. Convergence of parameter estimates was assessed through the visualisation of trace files in TRACER v1.7.2^73^ and the trees (excl. the first 15% of MCMC runs) were summarised with TREEANNOTATOR v1.10.4^74^.

### Characterisation of gene flow across Mediterranean *Puffinus*

To infer hybridisation events across all Mediterranean *Puffinus* populations, we first used Dtrios in Dsuite v0.5^30^ to calculate the sums of the three possible topologies (BABA, BBAA, and ABBA) and the resulting *D* and *f4* ratio statistics (Supplementary Table 11). To summarise the significant D-statistics comparisons and account for correlated results that may arise due to ancient gene flow, we performed the *f*-branch test (*f_b_*) implemented in Dsuite to assign gene flow to specific internal branches on the species tree. We identified significant instances of excess allele-sharing by setting a corrected p-value threshold using the Bonferroni correction, which was defined by dividing the p-value threshold of *p* < 0.01 by the number of cells in the *f*-branch matrix for which *f_b_* could be calculated. The results were visualised using the *dtools.py* script provided in the Dsuite package.

If gene flow is pervasive between a certain population pair, D-statistics lose their power to detect hybridisation events between their relative sister taxa, between whom hybridisation might still occur but at lower migration rates. In our case, the pervasiveness of hybridisation between the geographically closest populations (MAL, CAB and MEN) prevented us from detecting gene flow between PIT and MED, which are more isolated (Fig. 3a). To characterise whether hybridisation extended to these populations, we calculated the genomic proportion of the three topologies that included only the three main populations: PIT, MAL and MED. The underlying hypothesis was that if hybridisation or incomplete-lineage sorting (ILS) were scarce, the proportion of genome-wide topologies would be significantly skewed towards our species tree topology: ((PIT,MAL),MED). We used non-overlapping windows of 100 SNPs to build neighbour-joining trees with Phyml v3.3^82^ using a GTR model and the settings suggested by Martin and Van Belleghem (2017)^83^. Then we performed topology weighting in each of the 121,145 resulting trees with Twisst^83^ to determine the degree of monophyly for each topology.

Significance was assessed using Welch’s two-sample t-tests in R^84^ and corrected for multiple testing using Benjamini and Hochberg’s FDR-correction^85^.

To evaluate the patterns of hybridisation in MEN and CAB, the populations where gene-flow was suspected to be most pervasive, we attempted to take advantage of recent methods that use information about haplotype length and haplotype sharing patterns^86^. Such methods require the phasing of each diploid genome into two haploid sequences that reflect the linkage patterns across genomic variants^86^. We first performed a read-based phasing approach using WhatsHap v1.5^87^ for all individuals. The short insert sizes (350 bp) meant that such an approach would probably be unable to phase variants separated by distances greater than a few hundred base pairs. Therefore, we complemented the read-based phasing with the joint statistical phasing of all populations, performed using SHAPEIT4 v1.3^88^. The accuracy of the statistical phasing was assessed through the estimation of switch error rates when comparing with the read-based phasing using VCFtools v.0.1.15^59^ (option *--diff-switch-error*) (Supplementary Table 12).

The ChromoPainter and GLOBETROTTER pipeline has been shown to efficiently use haplotype information for accurate inferences of admixture^86,89,90^. The former software identifies the most probable source population for each of the haplotype blocks across an individual genome, while the latter uses its output to describe and date recent admixture events. To identify haplotype sharing patterns across all individuals, we ran ChromoPainter v2^89^ on the phased dataset with the adequate settings to paint all individuals by all others (*-a 0 0*). The switch rate (*-n 33700.840*) and global mutation probability (*-M 0.022739*) parameters were estimated in a previous run of ten Expectation-Maximisation (EM) algorithm iterations with a reduced subset of 16 individuals and 1,031,078 SNPs in accordance to the software guidelines^89^.

In order to date the admixture events that gave rise to the populations of MEN and CAB, we performed fastGLOBETROTTER^91^ runs using either MEN or CAB as the target population and the other three (PIT, MAL and MED) as source populations. We set the appropriate parameters to account for linkage patterns that might not be due to genuine admixture (*null.ind 1*) and we used shortened curve range (*curve.range: 0.5 5*) and bin width (*bin.width: 0.025*) parameters to accommodate the length of our scaffolds. We performed 100 bootstrap replicates for each of the fastGLOBETROTTER^91^ runs. Input files for fastGLOBETROTTER were generated using separate ChromoPainter v2^89^ runs for the generation of copy vector files and painting sample files (*-s 15*), as described in the fastGLOBETROTTER manual^91^. We visually checked that the fit of the modelled coancestry curves to the observed data supported the preferred hypothesis by fastGLOBETROTTER (Supplementary Fig. 19). For all estimates we used a generation time of 12.8 years^12^.

### Genome-wide scans for signatures of selection

Differentiation along the genome between Balearic and Yelkouan shearwaters was explored using a reduced subset of 24 individuals that excluded those most likely to result from recent introgression in CAB and MEN according to Struct-f4 (Supplementary Table 1). We estimated the relative differentiation between Balearic and Yelkouan shearwaters by calculating pairwise *F_ST_* in 25 kb non-overlapping sliding windows using the PopGenome v.2.7.5 R package^66^. Although *F_ST_* has been traditionally used to detect footprints of divergent selection, its interpretation can be misleading due to its susceptibility to a number of factors (incl. low recombination, demographic bottlenecks)^92,93^. In an attempt to overcome these caveats, we complemented our *F_ST_* genome scans with estimates of cross-population extended haplotype homozygosity (*XP-EHH*) averaged across the same 25 kb windows using the approach for unphased data implemented in selscan v2.0.0^94^. This haplotype-based method has proven useful to detect recent directional selection responsible for selective sweeps restricted to one of the populations by comparing haplotype lengths surrounding a given locus. Scaffolds that shared outlier windows (> 99.5% of all windows) for *F_ST_* and *XP-EHH* were identified as regions potentially under selection. Additionally, we also compared values for absolute (*d_xy_*) and net divergence (*d_a_*) between both taxa, as well as nucleotide diversity (*π*) within Balearic and Yelkouan shearwaters across the same 25 kb non-overlapping windows to deepen our understanding of the genomic landscape of differentiation.

Scans for the aforementioned statistics were repeated for scaffolds assigned to chromosome Z, with the exception of *XP-EHH*: the pervasiveness of missing data in this dataset precluded the confident use of selscan v2.0.0^94^ to calculate this statistic. In these scaffolds, window length was reduced to 15 kb to accommodate for the increased SNP density in Z chromosome scaffolds. The comparisons were performed between the two clusters of individuals identified in the PCAs of these sex-linked scaffolds, but individuals of unclear assignment to either group were excluded from the scans (Supplementary Table 1; Supplementary Fig. 20). In this case, we flagged contiguous outlier windows (> 99.5% of all windows) for *F_ST_* as potential targets of divergent selection between taxa. Finally, we evaluated syntenic genomic regions for all candidate autosomal and sex-linked scaffolds on closely-related species with chromosome-level genome assemblies (*Spheniscus humboldti* and *Ciconia maguari*) using minimap2 v2.11^63^ with default parameters.

### Characterisation of the main differentiation outlier between Balearic and Yelkouan shearwaters

Genomic footprints of linked selection and demography can be hard to distinguish and can lead to erroneous conclusions^92,93^. To disentangle the relative role of linked selection and demographic processes at driving differentiation, we focused on the study of the most representative autosomal *F_ST_* outlier region: scfF. To evaluate the pattern of differentiated SNPs across scfF, we performed local PCAs with the *lostruct* package in R^31^ using non-overlapping windows of 25 SNPs across this region. The distance between local PCAs was then calculated with the *pc_dist* function of the aforementioned package and visualised using multidimensional scaling (MDS) on two axes with the *cmdscale* function^31^. Furthermore, we quantified LD across these windows calculating pairwise LD as r^2^ using PLINK v1.90b5.3^64^. Finally, we visualised the distribution and extent of haplotypes within this region across the sampled individuals with GenotypePlot (https://github.com/JimWhiting91/genotype_plot) in R. The same visualisation was repeated for each of the *F_ST_* and *XP-EHH* shared outliers: none showed haplotype segregation akin to that observed in scfF, reinforcing our confidence in the choice of scfF to study divergent selection between Balearic and Yelkouan shearwaters (Supplementary Fig. 21).

We identified genes present within scfF (as well as all other windows flagged as outliers) by referring to the Balearic shearwater annotated genome^19^. These genes might potentially be under divergent selection between these taxa; their function and role in adaptation in other organisms was characterised through the available bibliography (Supplementary Table 6). We also identified non-synonymous SNPs whose genotype across individuals coincided with the distribution of scfF haplotypes (Supplementary Table 1). The evolutionary conservation of the coding regions within each gene, which has been suggested to be indicative of their functional constraint^95^, was later evaluated by comparing phyloP scores across a 364-way bird genome alignment (see “Estimates of genetic load using evolutionary conservation”).

### Investigating the role of balancing selection and migration in adaptation

Selection on ancient genetic modules maintained through balancing selection can be an effective method for rapid local adaptation^37^. In our case, we wondered whether the intermediate scfF haplotype frequencies in Balearic shearwater could be indicative of such a process. We first investigated this possibility by calculating the allele frequency correlation summary statistic, ß, in 25 kb sliding windows (*-w 25000*) as implemented in BetaScan2^96^ using all Balearic individuals excluding CAB (*n=17*). High values of this statistic reflect clusters of polymorphisms under similar allele frequencies^96^. By setting the minimum core allele frequency (*-m*) at *m=0.25* we evaluated whether these clusters were found at intermediate allele frequencies, a characteristic of regions under balancing selection. These values were later averaged across 25 kb non-overlapping windows. Additionally, we calculated Tajima’s D values across the same windows with PopGenome^66^ to infer deviations from the neutral SFS expectations. Comparison between scfF and genome-wide values for both statistics were performed using Mann-Whitney U-tests in R^84^.

Gene flow can result in similar haplotype frequencies as those explained by balancing selection, and both processes might be difficult to differentiate in the light of pervasive hybridisation^38,97^. We attempted to differentiate these processes by building neighbour-joining genealogies between the phased haplotypes of HOM1 individuals of both *Puffinus* using 100 SNP non-overlapping windows with Phyml^82^ (see “Characterisation of gene flow across Mediterranean *Puffinus*”). If ancestral balancing selection were driving intermediate haplotype frequencies in Balearic shearwater, H1 haplotypes from Balearic shearwater should form a clade that would have diverged from Yelkouan shearwater during the speciation process. Otherwise, multiple recent coalescent events between H1 haplotypes of Balearic and Yelkouan shearwaters would indicate the recent introduction of H1 haplotypes into Balearic populations via hybridisation.

Moreover, genomic windows under positive selection are likely to show signatures of reduced recombination rates, as the breakdown of adaptive haplotypes through recombination would have negative effects on an individuals’ fitness. To test whether scfF fulfilled this assumption in homozygous Yelkouan shearwaters (*n=10*), we performed two estimates of recombination across 25 kb windows with PopGenome^66^: the average pairwise r^2^ value (*ZnS*) as a measure of LD, and the scores for the 4-gamete test. We compared these values with two sets of Balearic shearwater individuals: HOM0 (*n=8*) and HOM1 (*n=3*) individuals for scfF (Supplementary Table 1). To control for the effect of different sample sizes per group, we obtained Z-scores for *ZnS* and 4-gamete test results by normalising them with values across all other 1 Mb regions syntenic with chromosome ends in both *S. humboldti* and *C. maguari*, as identified with minimap2 v2.11^63^ (Supplementary Table 13). Significance of differences between groups was assessed using non-parametric Mann-Whitney *U*-tests in R^84^ and corrected for multiple testing using Benjamini and Hochberg’s FDR-correction^85^.

### Genomic diversity and inbreeding

To assess the potential loss in genetic diversity derived from population declines in Mediterranean *Puffinus,* we first calculated genome-wide heterozygosity for each individual. Due to the sensitivity of individual-based estimates of heterozygosity to sequence coverage, we generated individual VCF files from the raw autosomal VCF and kept only SNPs with depths between 0.5x and 2x the individuals’ average coverage (Supplementary Table 10). We then obtained individual heterozygosity estimates by calculating the number of heterozygous positions per base pair along 25 kb non-overlapping autosomal windows with VCFtools v.0.1.15^59^, subtracting the masked proportion of each window from each denominator. These were later averaged to obtain genome-wide average heterozygosities for each individual.

We also identified runs of homozygosity (ROH) for each individual genome using PLINK v1.90b5.3^64^ and later estimated individual inbreeding coefficients (*F_ROH_*) as the proportion of each genome occupied by ROH. For the identification of ROH with PLINK^64^, we used a scanning window size of 50 SNPs (*--homozyg-window-snp* 50) and kept the default settings for minimum SNP density (--*homozyg-density*), maximum gap between SNPs (--*homozyg-gap*) and number of heterozygous sites within ROH (--*homozyg-het*) because they did not affect the identification of ROH^98^. Due to the low coverage of our data, we tolerated heterozygous and/or missing sites that might have been derived from sequencing errors with the following parameters: we allowed up to five heterozygous SNPs per scanning window (*-- homozyg-window-het 5*) and up to 50 SNPs with missing data (*--homozyg-window-missing 50*). Additionally, we ensured the proper delimitation of ROH edges including SNPs that were present in at least 10% of the scanning windows of a ROH (*--homozyg-window-threshold 0.1*). We divided ROH into two categories according to their length: short ROH (100 kb - 1 Mb) and long ROH (> 1 Mb). Finally, we removed all identified ROH with a high proportion of heterozygous sites (*PHET* > 0.03); this threshold was chosen as it was empirically shown to avoid the inclusion of low-diversity regions resulting from selective sweeps. We evaluated the significance of all pairwise inter-population comparisons for average genome-wide heterozygosity and *F_ROH_* using non-parametric Mann-Whitney *U*-tests in R^84^. We corrected p-values for multiple testing using Benjamini and Hochberg’s FDR-correction^85^.

### Estimates of genetic load using evolutionary conservation

To estimate proxies of evolutionary conservation, we added the Balearic shearwater reference genome to the 363-way genome alignment generated by the B10K project^95^ by adding a new branch sister to the Cory’s shearwater (*C. borealis*). We followed the strategy described in Armstrong *et al.* (2020)^99^ by running two separate alignments using Progressive Cactus and updating the 363-way alignment with *halAddToBranch*. We then used PhyloP^39^ to calculate per-base constraint and acceleration p-values on the macrochromosomes of the Balearic shearwater-referenced 364-way alignment. We calculated PhyloP scores under the null hypothesis of the neutral evolution model for macrochromosomes presented in Feng *et al.* (2020)^95^ using a likelihood ratio test at each alignment column (*--method LRT*) with only constraint scores output (*--mode CON*). Afterwards, we converted phyloP scores (which represent log-scaled p-values of conservation) into p-values, and subsequently into q-values using the FDR-correction of Benjamini and Hochberg.

We observed that the q-value of a given SNP depended on its frequency within a population: decreasing q-values tended to show low minor allele frequencies within a population’s folded SFS, as would be expected with increasing deleteriousness (Supplementary Fig. 22). Consequently, we used a stringent q-value threshold of 1×10^-8^ to delimit a highly phylogenetically conserved dataset of 6,134 SNPs that most probably contribute to genetic load within *Puffinus* populations. To ensure that the polarisation of the variants reflected their deleteriousness, we extracted the base composition of each significantly conserved position in the 364-way alignment using *halSnps* and polarised these SNPs with the major allele across the avian phylogeny using a custom script. Finally, we also removed all those SNPs whose derived allele was also present in at least one individual of Manx shearwater; these SNPs likely represent either wrong polarisations or positions that are adaptive across most North Atlantic *Puffinus* species. With this filtered dataset, we used the number of deleterious variants in homozygosity and heterozygosity as proxies for realised and masked load, respectively. Furthermore, we also counted the number of shared deleterious alleles between all pairs of individuals to understand the extent of common load within and between populations/taxa. We evaluated the significance of all pairwise inter-population comparisons through the use of non-parametric Mann-Whitney *U*-tests in R^84^. We corrected p-values for multiple testing using Benjamini and Hochberg’s FDR-correction^85^.

### Estimates of genetic load in coding regions

In order to assess mutational load in coding regions of the shearwater’s genome, we annotated our autosomal SNP dataset and predicted their functional effects using SnpEff v5.1^40^. Previously, we generated a SnpEff database for the Balearic shearwater using the annotated reference genome of the species. To ensure that potential annotation errors would not bias our load estimates, we used a high confidence gene set including those genes annotated in the Balearic shearwater reference genome that had homologs in either the Cory’s shearwater, zebra finch or human genomes. As the identification of non-synonymous variants depends on which allele is considered as derived, we also polarised all alleles using the major allele across the 364-way cactus alignment. Furthermore, we removed positions where the derived allele was present in at least one individual of Manx shearwater to minimise polarisation errors. Variants were separated into the four impact categories defined by SnpEff v5.1^40^: “High”, “Moderate”, “Low” and “Modifier”. While the first three categories aim to represent a decreasing functional effect of the variants (from nonsense variants to synonymous mutations), “Modifier” variants, usually placed in non-coding regions, are generally assumed to be neutral although their effect is difficult to predict^40^. Therefore, the number and genotype (homozygous/heterozygous) of variants in each individual was inferred for the first three categories. As with evolutionarily conserved SNPs, we also counted the number of deleterious alleles shared between individual pairs. As normality was fulfilled for all categories, the significance of pairwise inter-population comparisons was evaluated using Welch’s two-sample t-tests in R^84^ and p-values were corrected for multiple testing using Benjamini and Hochberg’s FDR-correction^85^.

### Forward simulations

To understand the potential consequences of inbreeding and hybridisation in the remaining population of Balearic shearwaters under current demographic projections, we performed individual-based forward simulations with SLiM v4.0.1^23^ using its non-Wright-Fisher (non-WF) implementation. The use of non-WF models allows the construction of models with fully customisable life-history traits, which we took advantage of to build simulations with overlapping generations and realistic age-class distributions, age-based mortality and age-based reproductive probabilities obtained from previous viability analyses^12^ (Supplementary Tables 8 & 9). In order to explore the evolution of population size, heterozygosity and ROH in the future, we simulated 25 Gbp recombining DNA fragments for two populations representing Balearic and Yelkouan shearwaters and ran the different models for 500 cycles. Recombination rate was set at a uniform rate of *r* = 1×10^-8^ crossovers/bp/generation. In all cases, mutations were introduced to the output tree sequences after the completion of simulations using msprime v.1.2.0^81^.

To use populations that are accurate in terms of diversity and differentiation as the starting-point for our simulations, we ran two burn-in steps to achieve current estimates of heterozygosity and *F_ST_* based on the results of the current study. First, we simulated a single panmictic population with a carrying capacity of *K_0_* = 88,000 individuals using a scaled-up mutation rate of 0.758×10^-8^ mutations/bp/generation until a heterozygosity of 0.0019 bp^-1^, which happened after 180,900 cycles. Next, we split this population into two subpopulations with initial carrying capacities comparable to current census estimates for Balearic (*K_1_* = 10,000 individuals) and Yelkouan shearwaters (*K_2_* = 88,000 individuals). Simulations were run until *F_ST_* between the two populations reached values similar to the ones observed in this study (*F_ST_* = 0.017) using the same scaled-up mutation rate as before. Once this value was reached after 1,400 further cycles, we output the entire genealogy via tree-sequence recording and used it as input for each of the future models.

We simulated nine distinct scenarios based on the combination of survival/reproductive probabilities and migration rates. On the one hand, we compared the outcome of three possible unidirectional migration scenarios: a) no hybridisation, potentially due to the local extinctions of MEN and CAB; b) current migration rates into Balearic populations (*m = 10 migrants/generation*); and c) potential human-mediated translocation (*m = 100 migrants/generation*). In all cases, migrants were randomly drawn from a uniform distribution of age classes (3-49 years). On the other hand, we compared the results under three demographic scenarios using life-history parameter estimates outlined in a recent viability analysis^12^: a) current demographic projections; b) demographic projections assuming increased survival probabilities due to immediate bycatch reduction; c) projections if the implementation of bycatch reduction measures is delayed 25 years. For the first two models, survival and reproductive probabilities for each age class were simplified from the deterministic models presented in Genovart *et al.* (2016)^12^ (Supplementary Table 9; Supplementary Fig. 23 & 24). For the latter model, the first 25 cycles used current parameter estimates (as in model A) but the following cycles used the reduced bycatch estimates (model B). Maximum longevity was set at 75 years in all simulations. While population sizes in Balearic shearwaters varied according to the survival/reproductive probabilities set by each model, the Yelkouan population remained stable around its carrying capacity of *K_2_* = 88,000 individuals. All simulations were run for a total of 500 cycles and used a mutation rate of 0.758×10^-8^ mutations/bp/generation.

We kept track of population size at each cycle in SLiM and repeated the simulations five times per model to obtain confidence intervals. Furthermore, we output VCFs with ten simulated individuals at six points in time (*t* = (0, 25, 50, 100, 250, 500)) for each simulation using tskit v0.5.6 to evaluate the evolution of heterozygosity and *F_ROH_*. To calculate these two parameters, we used the same procedure as the one we employed on resequenced genomes, employing VCFtools and PLINK (see “*Genomic diversity and inbreeding*”). We evaluated the significance of pairwise comparisons between models and time points using non-parametric Mann-Whitney *U*-tests in R^84^. We corrected p-values for multiple testing using Benjamini and Hochberg’s FDR-correction^85^.

## DATA AVAILABILITY

Raw reads generated for this study are archived on the European Nucleotide Archive (ENA) under accession number PRJEB75355.

## CODE AVAILABILITY

All code written for this project is available on the following GitHub repository: https://github.com/molevol-ub/Puffinus_mauretanicus_popgen.

## ACKNOWLEDGEMENTS

We are grateful to Meritxell Genovart, Daniel Oro, Paulo Lago and Martin Austad for kindly providing samples. We would like to thank Alejandro Sánchez-Gracia for expert advice on analyses and Martí Franch for his wonderful shearwater illustrations. Permits required for conducting this study were provided by the following local or national authorities (permit references in brackets): Conselleria de Medi Ambient del Govern Balear; Environment & Resources Authority from Malta (NP0003/18); Istituto Superiore per la Protezione e la Ricerca Ambientale – San Domino, Tavolara/Molara, Cavoli (Prot. 37088, 21/06/2016); Direzione Generale della Difesa dell’Ambiente, Regione Autonoma della Sardegna – Tavolara/Molara, Cavoli (Determinazione Prot 12498, Rep 255, 28/06/2016); Dipartemento Regionale dello Sviluppo Rurale e Territoriale, Regione Siciliana – Lampedusa (Prot 11382, 09/05/2016); Ministarstvo Zastite Okolisa i Prirode, Republika Hrvatska - Lastovo (klasa UP/I-612-07/16-48/20, 10/03/2016; Agence de Protection et de l’Aménagement du Littoral; and Armée tunisienne (2014). This study was carried out with partial financial support from the Fundación

Banco Bilbao Vizcaya Argentaria (Project 062-17), and from the Ministerio de Economía y Competitividad of Spain (projects CGL2016-78530-R, PGC2018-093924-B-100, and PID2019-103947GB).

