## Supplementary Figures 1-24 for "Pervasive and recurrent hybridisation prevents inbreeding depression in Europe’s most threatened seabird"

-----

### Supplementary Figures

Guillem Izquierdo-Arànega[1,2], Cristian Cuevas-Caballé[2,3], Francesco Giannelli[4], Josephine Rosanna Paris[4], Karen Bourgeois[5], Emiliano Trucchi[4], Jacob González-Solís[1,2], Marta Riutort[2,3], Joan Ferrer Obiol[6]* & Julio Rozas[2,3]*

[1]Departament de Biologia Evolutiva, Ecologia i Ciències Ambientals, Universitat de Barcelona, Barcelona, Spain

[2]Institut de Recerca de la Biodiversitat, Universitat de Barcelona, Barcelona, Spain

[3]Departament de Genètica, Microbiologia i Estadística, Universitat de Barcelona, Barcelona, Spain

[4]Department of Life and Environmental Sciences, Marche Polytechnic University, Ancona, Italy

[5]Aix Marseille Université, CNRS, IRD, Avignon Université, Institut Méditerranéen de Biodiversité et d'Ecologie marine et continentale, Bât. Villemin, Technopôle Arbois-Méditerranée, UMR IMBE, Aix-en-Provence, France

[6]Dipartimento di Scienze e Politiche Ambientali, Università degli Studi di Milano, Milano, Italy

* *Co-corresponding authors.*

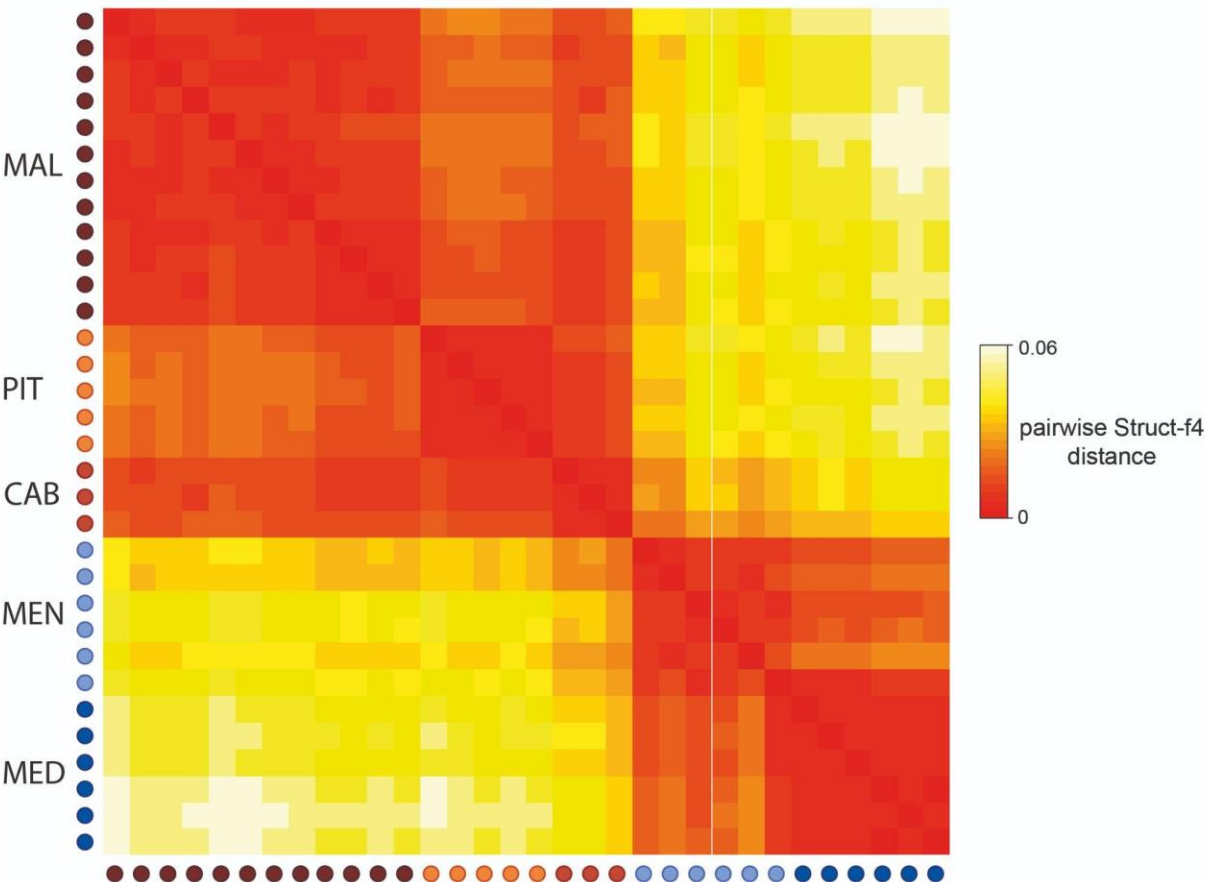


**Supplementary Fig. 1:** Pairwise distance matrix of Struct-f4[1] genetic distances between sampled individuals. Increasing genetic affinities are indicated by a yellow-to-red gradient.

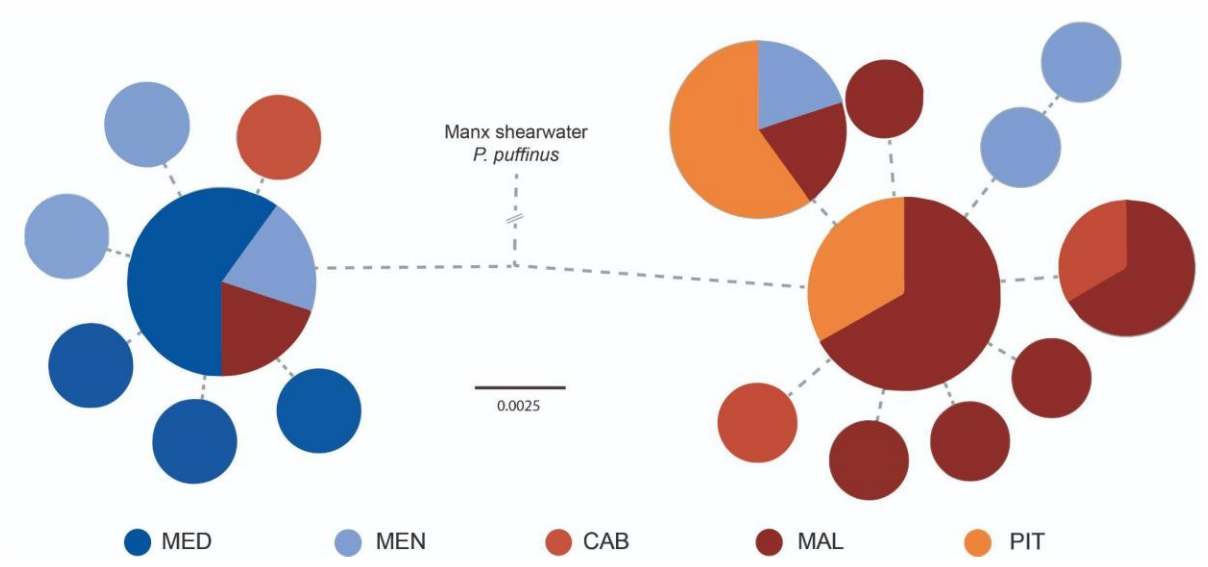


**Supplementary Fig. 2:** Haplotype genealogy graph obtained from whole mitochondrial genomes of Mediterranean *Puffinus* using 393 mitochondrial SNPs; minimum edge length was set to $e = 3$.

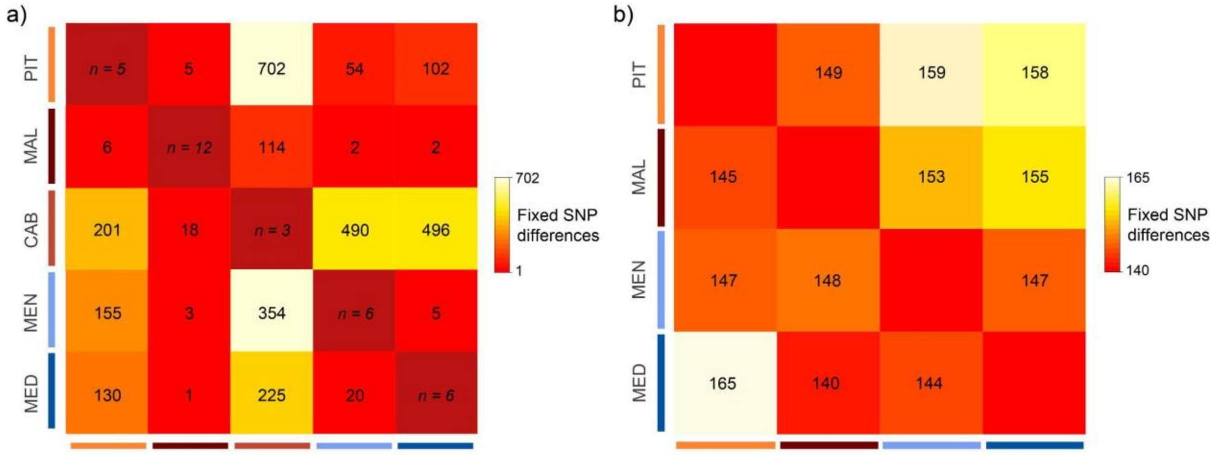


**Supplementary Fig. 3:** Colour-coded heatmaps according to the number of fixed SNPs between population pairs using: all individuals from each population **(a)**; or five individuals from all populations except CAB **(b)**. In **(a)** the number of individuals per population is indicated across the diagonal. In both cases, values above the diagonal show fixed differences in autosomal scaffolds, while values below the diagonal show those within sex-linked scaffolds.

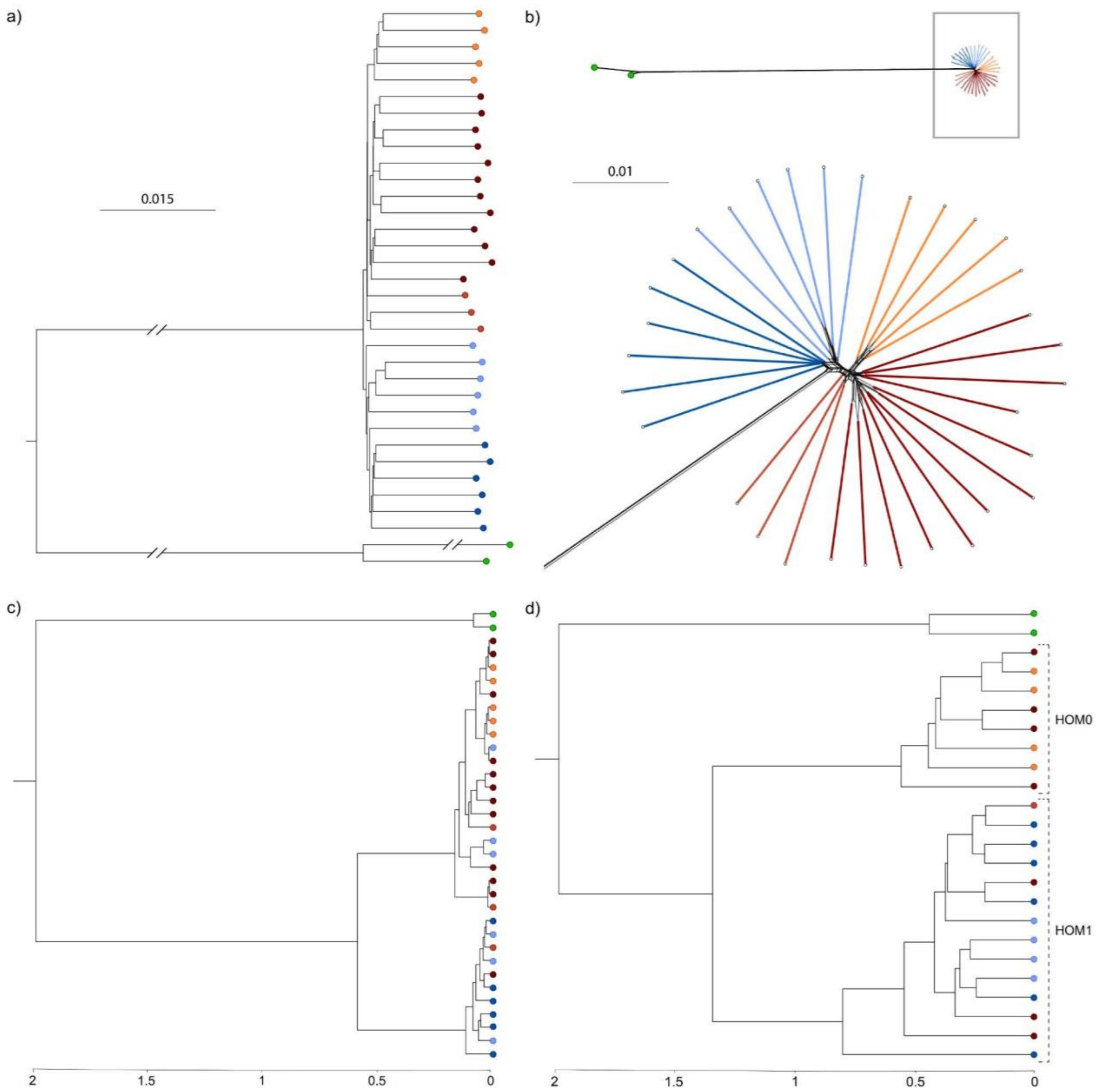


**Supplementary Fig. 4: a,** Neighbour-joining phylogenetic tree inferred with the ape[2] R package using 1,304,832 autosomal SNPs. **b,** Reticulation network inferred using the Neighbour Net agglomerative algorithm[3] based on pairwise distances using 1,304,832 autosomal SNPs. **c,d,** Dated multispecies coalescent trees obtained using: whole mitochondrial genome sequences **(c)**; and 603 SNPs found within the centre of the selective sweep in scfF **(d).** Each terminal corresponds to one individual, whose source population is indicated by the colour of the tip as coded in Fig.1.

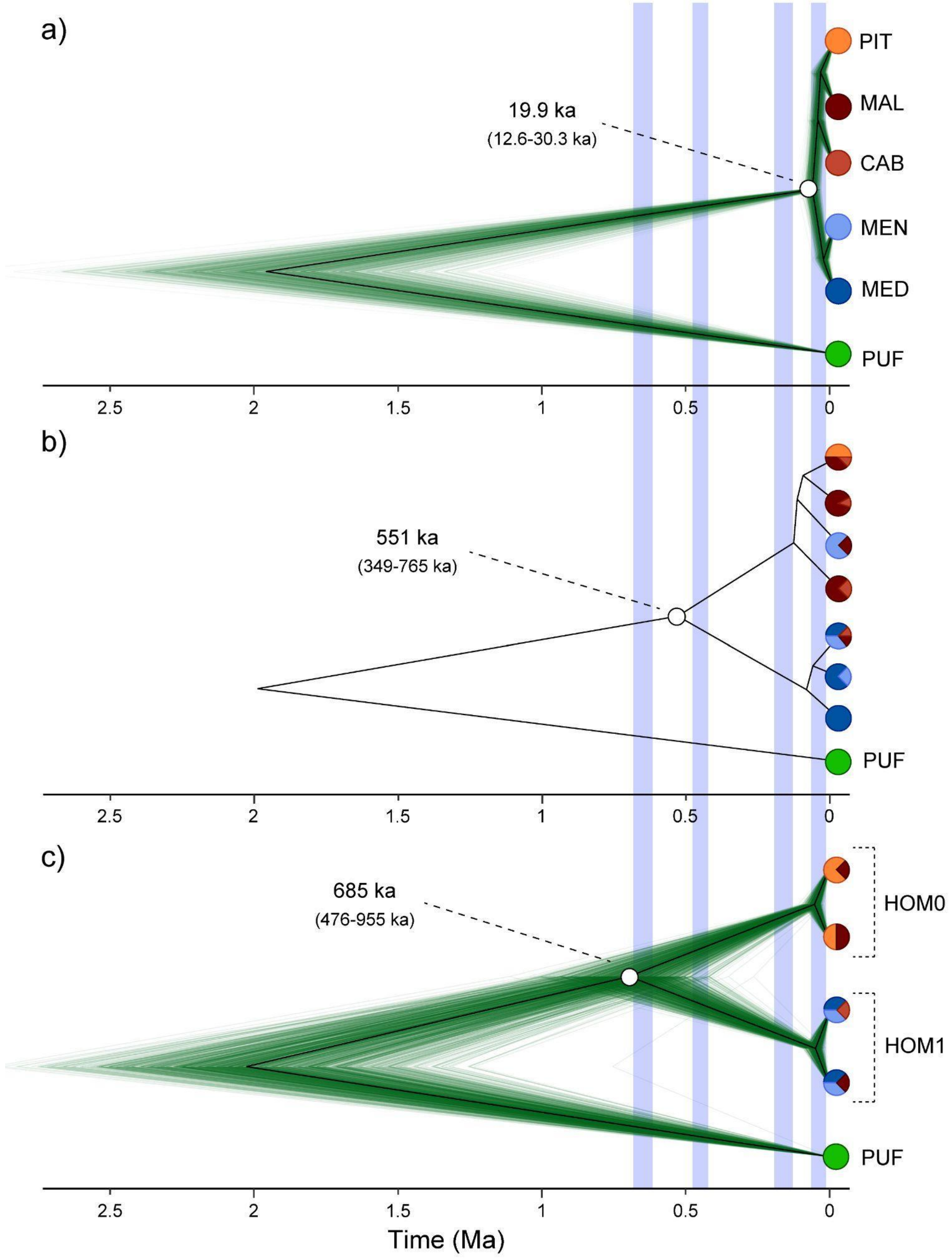


**Supplementary Figure 5: a-c,** Dated multispecies coalescent trees using 5000 random genome-wide SNPs **(a);** whole mitochondrial genome sequences **(b)**; and 603 SNPs found within the centre of the selective sweep in scfF **(c)** (see *Genome-wide scans for signatures of divergent selection*). In the latter, heterozygous individuals for the two main haplotypes were not included. Pie charts represent the proportion of individuals of each population in any given clade. In **a)** and **c)**, thin green lines show individual trees sampled from the posterior distribution; the black line indicates the maximum-clade-credibility summary tree. Blue bars highlight the main Pleistocene glacial maxima.

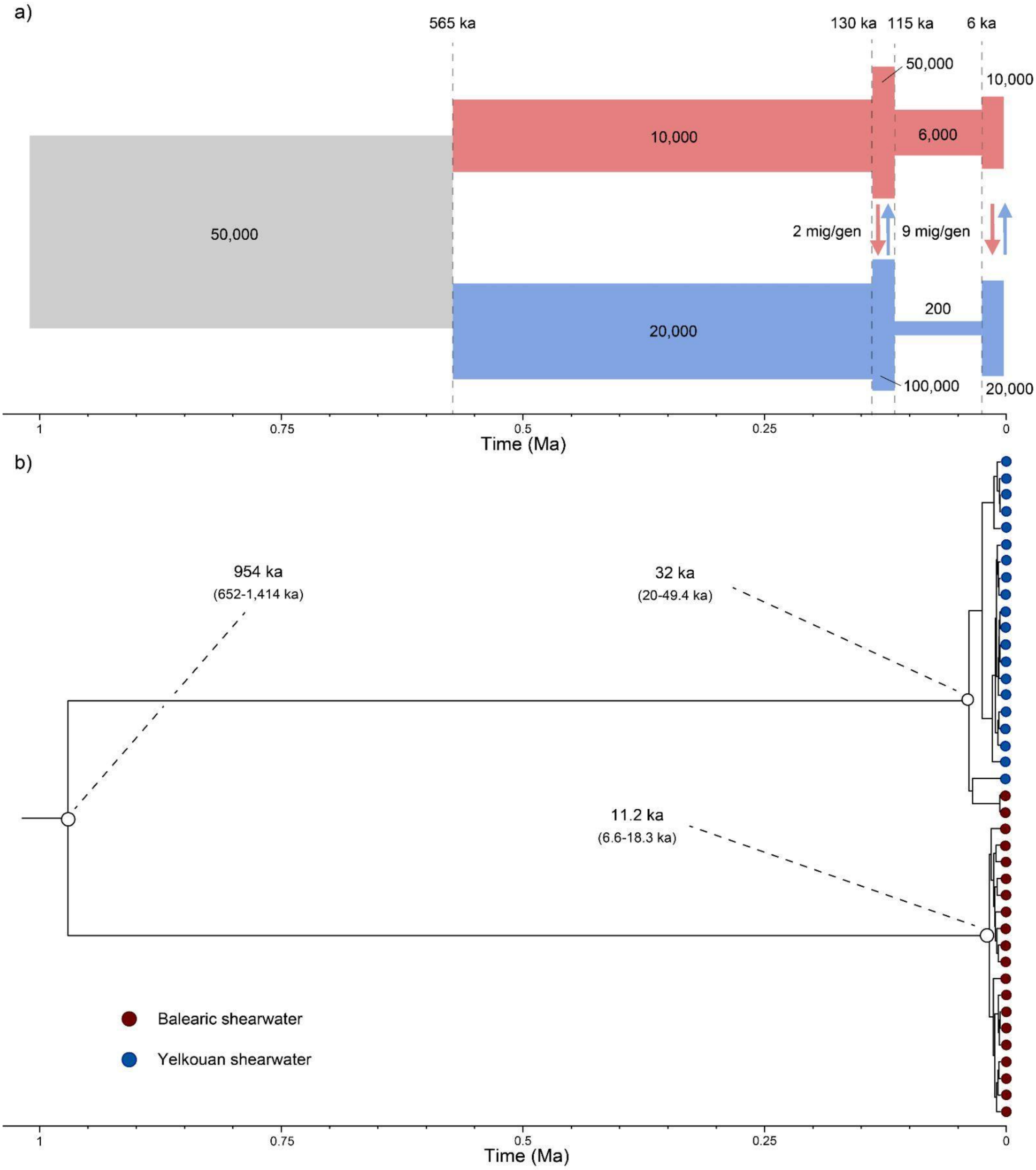


**Supplementary Fig. 6: a,** Schematic representation of the evolutionary model for which we simulated whole mitogenomes in SLIM[4]. Time is indicated in thousands of years (ka), population size in number of individuals and migration rates in number of migrants per generation. **b,** Dated multispecies coalescent trees using whole simulated mitogenomes in

SLiM[4] under the scenario illustrated in **(a)**. Coloured tips indicate the taxonomic assignment of each of the simulated individuals.

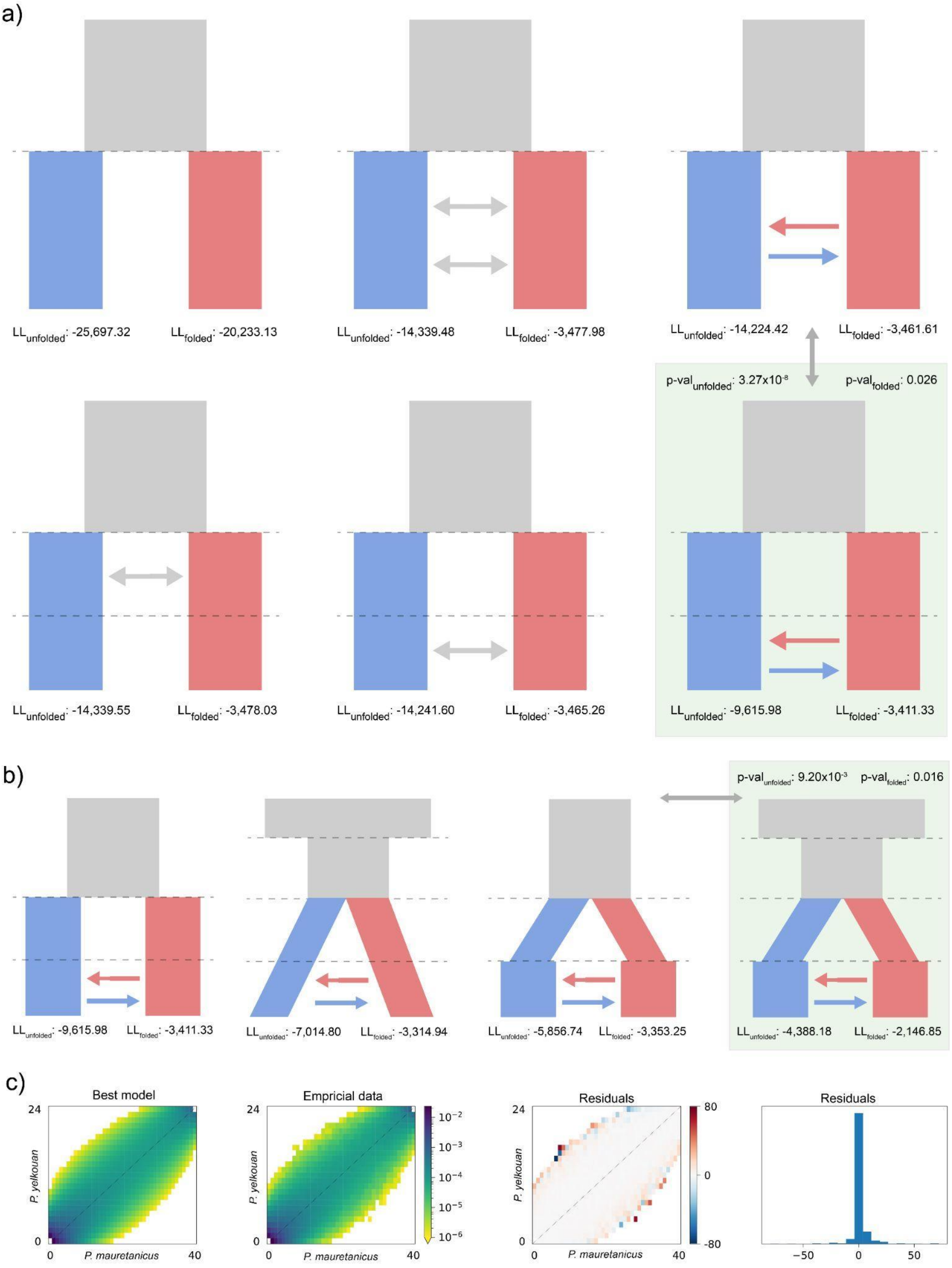


**Supplementary Fig. 7: a,b,** Demographic models compared in ∂a∂i[5] with the 2D-SFS of Balearic and Yelkouan shearwaters. In **(a)** we show models that ignored demographic size changes; the best such model was compared with models shown in **(b)**, which take population size oscillations into account. Log-likelihoods for all models using both the folded and unfolded 2D-SFS are shown, as well as the significance of comparisons between the best model for each **(a)** and **(b)**, highlighted in green, with the next best model (connected by grey arrows). **c,** Comparison of the unfolded 2D-SFS of the best-fitting model (left), our data (centre), and the resulting residuals (right). In the residual plot, red colours show an excess of alleles at a certain frequency in the empirical 2D-SFS compared to the best model, while blue show a deficit.

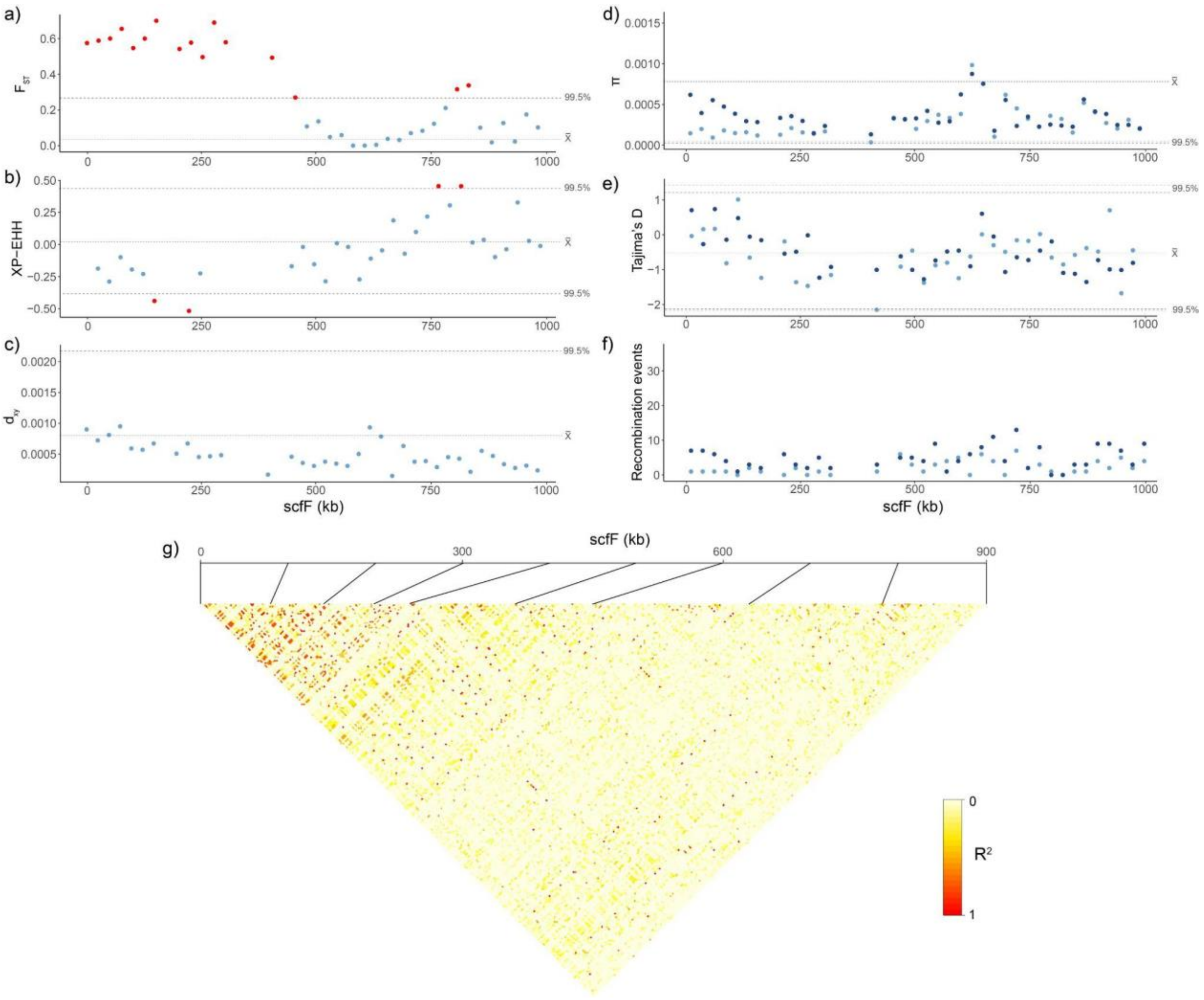


**Supplementary Fig. 8: a-f,** Genome scans using 25 kb non-overlapping sliding windows across the most likely candidate window for divergent selection between Balearic and Yelkouan shearwaters - scfF - for: $F_{ST}$ **(a)**, XP-EHH **(b)**, $d_{xy}$ **(c),** $\pi$ **(d),** Tajima's D **(e)** and 4-gamete test results **(f).** In the case of $\pi$, Tajima's D and 4-gamete test results, dark dots indicate values for Balearic and light dots for Yelkouan shearwaters. Dotted lines, where present, indicate average genome-wide values and 99.5% outlier thresholds. **g,** Pairwise linkage disequilibrium (LD) heatmap for scfF calculated as $r^2$.

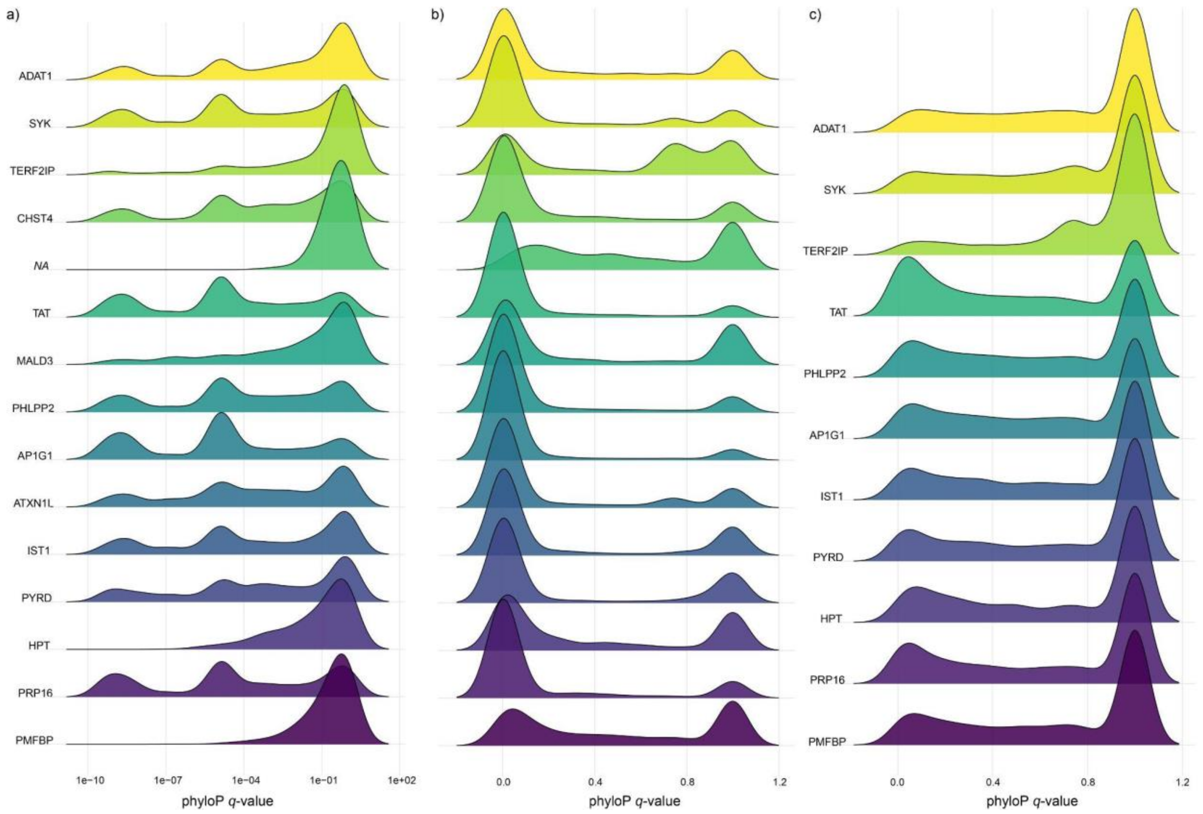


**Supplementary Fig. 9:** Density plots of phyloP[6] *q*-values for positions found within the coding **(a,b)** and intronic regions **(c)** of genes found within scfF. Smaller *q*-values represent sites that are under a heavier evolutionary constraint. Plots of sites in coding regions are shown over a log-transformed **(a)** or untransformed X axis **(b)**.

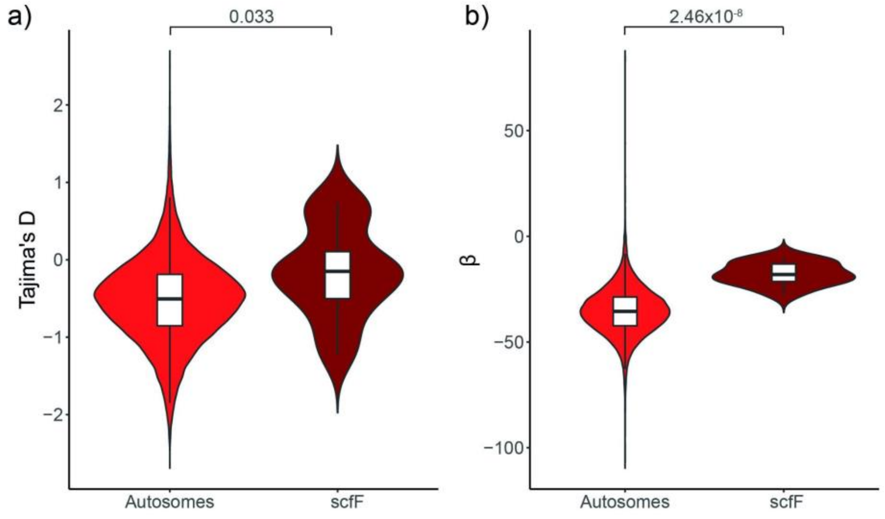


**Supplementary Fig. 10:** Comparison of summary statistics and neutrality tests between 25 kb windows within scfF and across the rest of the genome in Balearic shearwaters (excluding CAB): Tajima's *D* **(a)** and *ß* **(B)**. P-values are shown for significant pairwise comparisons (Mann-Whitney *U*-test).

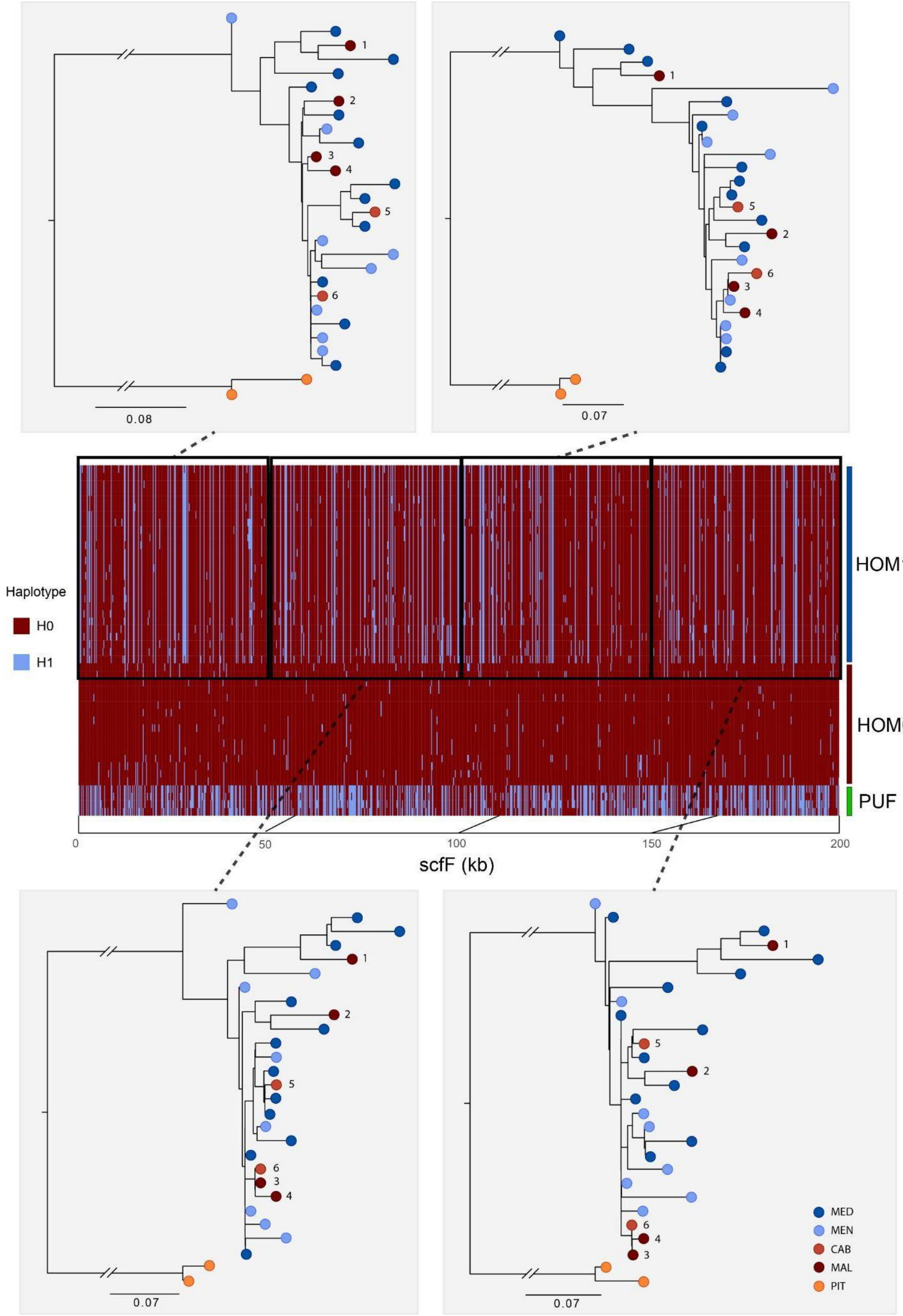


**Supplementary Fig. 11:** Phased haplotype plot of homozygous individuals for haplotypes within scfF polarised for the major allele in the Balearic shearwater. Reference SNPs (H0) are marked in dark red and alternate SNPs (H1) in blue. Neighbour-joining genealogies for each non-overlapping 100-SNP window found within scfF are shown, with tips colour-coded according to its population as indicated in Fig. 1. H1 haplotypes belonging to Balearic shearwaters are identified with the same number across all trees.

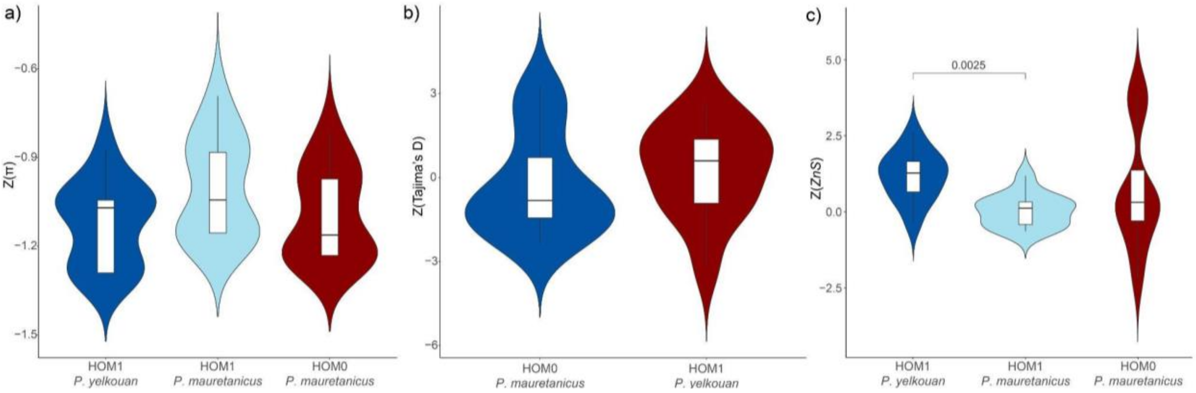


**Supplementary Fig. 12: a,** Normalised genetic diversity (*π*) measured across nonoverlapping 25kb windows within scfF in HOM0 and HOM1 individuals. **b,** Normalised Tajima's D values across nonoverlapping 25 kb windows within scfF in HOM0 and HOM1 individuals. **c,** Recombination rates of H0 and H1 scfF haplotypes in Balearic and Yelkouan shearwaters, measured as normalised *ZnS* values across 25 kb windows. In all cases, FDR-corrected p-values are shown for significant pairwise comparisons (Mann-Whitney *U*-test).

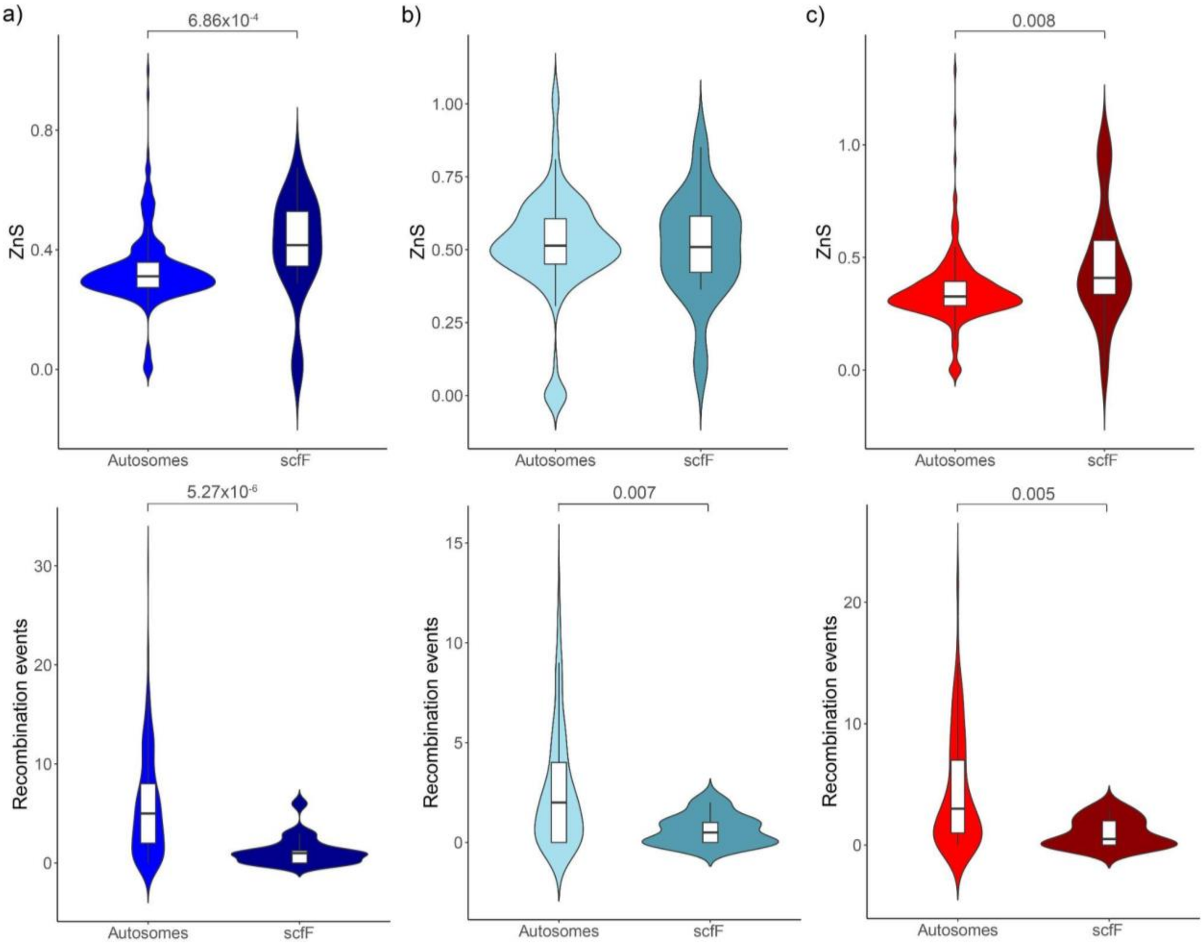


**Supplementary Fig. 13: a-c,** Comparison of recombination rates between scfF and other chromosome ends in: Yelkouan shearwaters **(a)**; HOM1 Balearic shearwaters **(b)**; and HOM0 Balearic shearwaters **(c)**. Values are shown for ZnS (top) and 4-gamete test scores (bottom) calculated across 25 kb windows. In all cases, p-values are shown for significant pairwise comparisons (Mann-Whitney *U*-test).

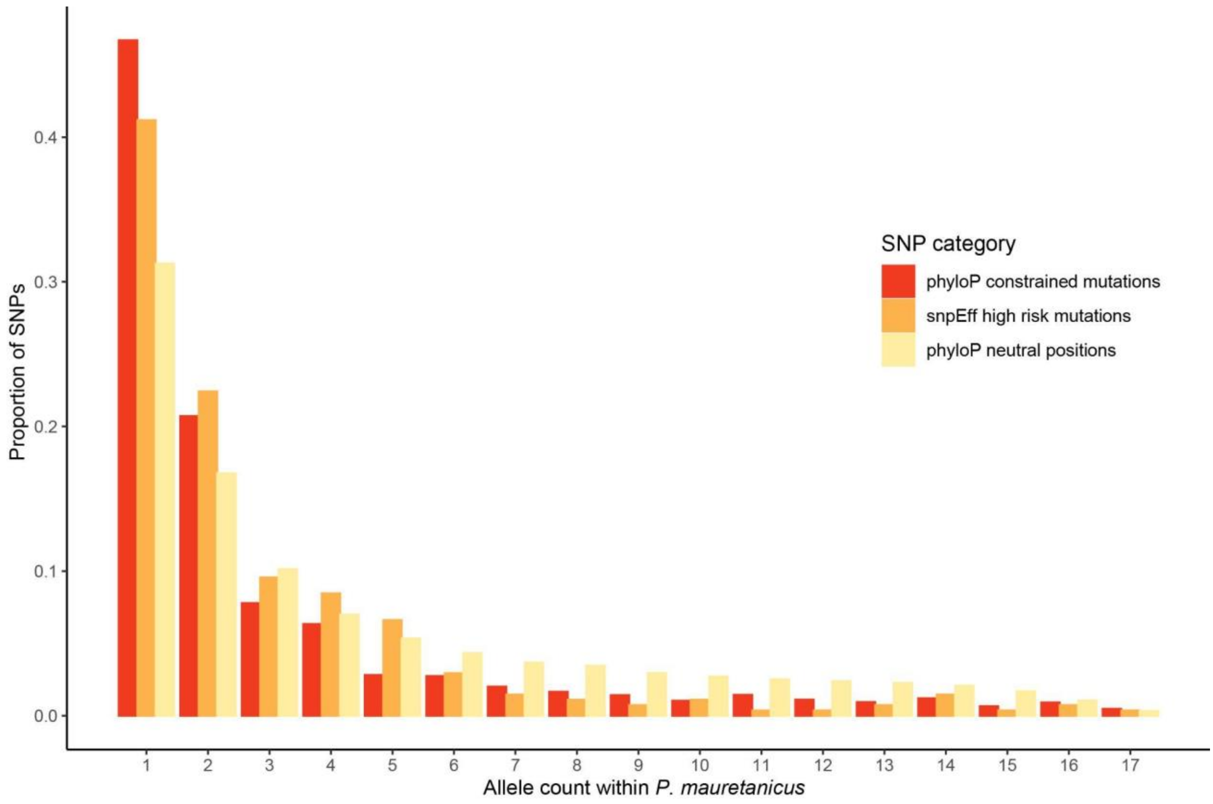


**Supplementary Fig. 14:** Folded site frequency spectrum (SFS) of Balearic shearwaters (PIT and MAL) using three different SNP datasets: potentially neutral SNPs (phyloP score = 0) in yellow; high risk SNPs found within coding regions as inferred by snpEff[7] in orange; and evolutionary constrained SNPs inferred with phyloP[6] in dark red.

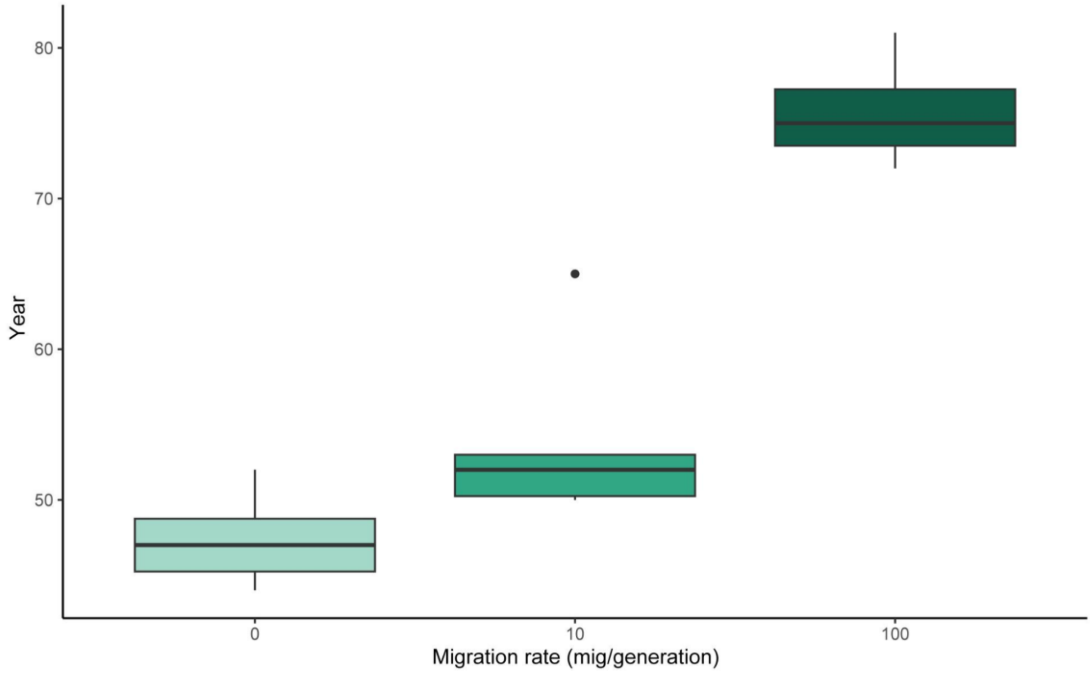


**Supplementary Fig. 15:** Predicted year of virtual extinction (N < 10 individuals) for the Balearic shearwater across simulations using current demographic projections[8] and different migration rates with the Yelkouan shearwater. Boxplots show the median (centre), the first and third quartiles (box bounds), the smallest and largest values within 1.5x the interquartile range from the first and third quartiles (whiskers), and outliers (points).

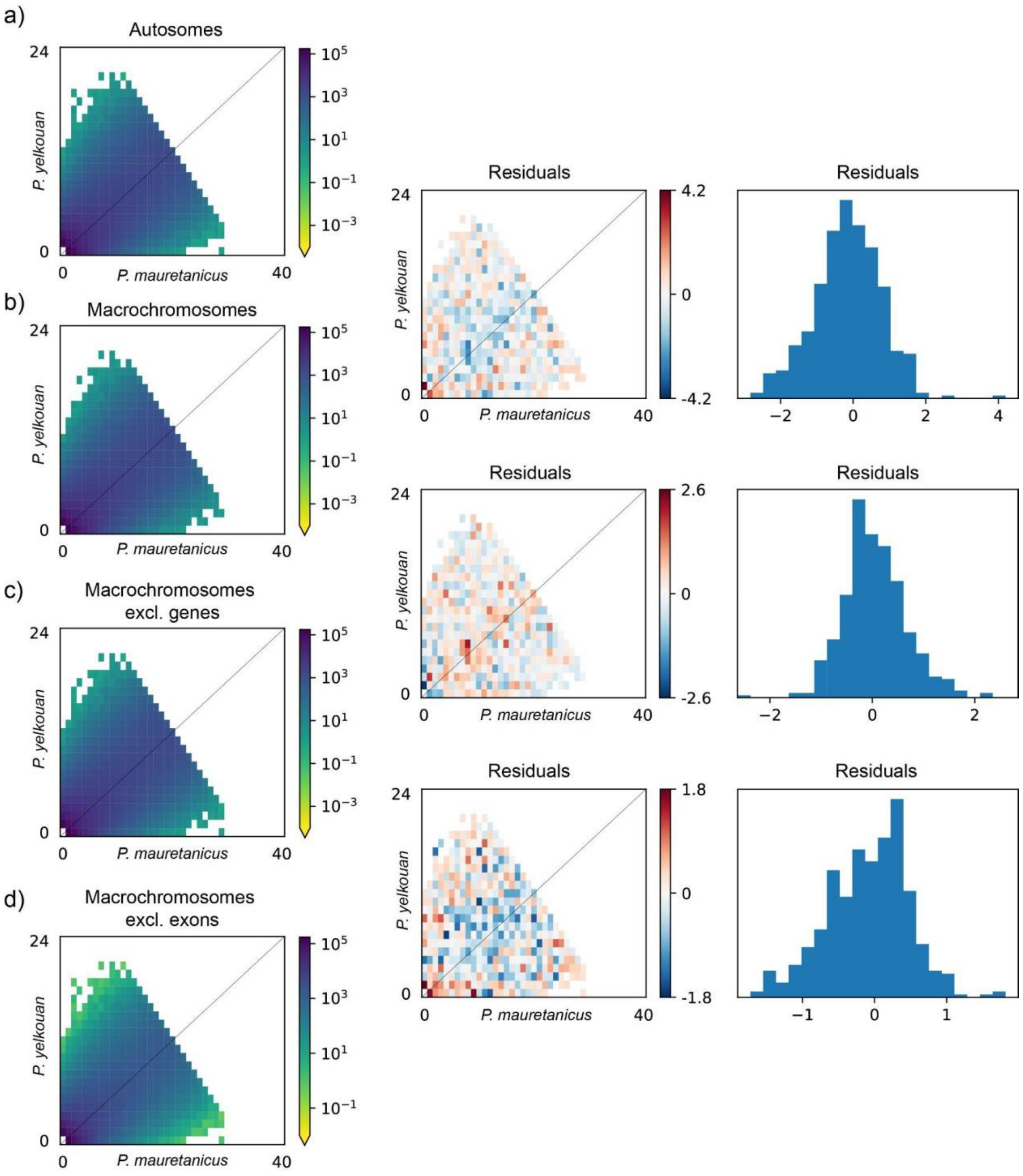


**Supplementary Fig. 16:** Comparison of the folded 2D-SFS of Mediterranean *Puffinus* using several SNP datasets: all 1,090,933 autosomal SNPs **(a)**; 859,300 SNPs syntenic with macrochromosomes in *Spheniscus humboldti* **(b)**; 685,502 SNPs found within macrochromosomes, excluding genes **(c)**; 848,800 SNPs found in macrochromosomes outside coding regions **(d)**. Residuals from the comparison between each dataset and the preceding one in number of SNPs are shown next to both models being compared. Red colours show a deficit of alleles at a certain frequency compared to the preceding dataset, while blue show an excess of said alleles.

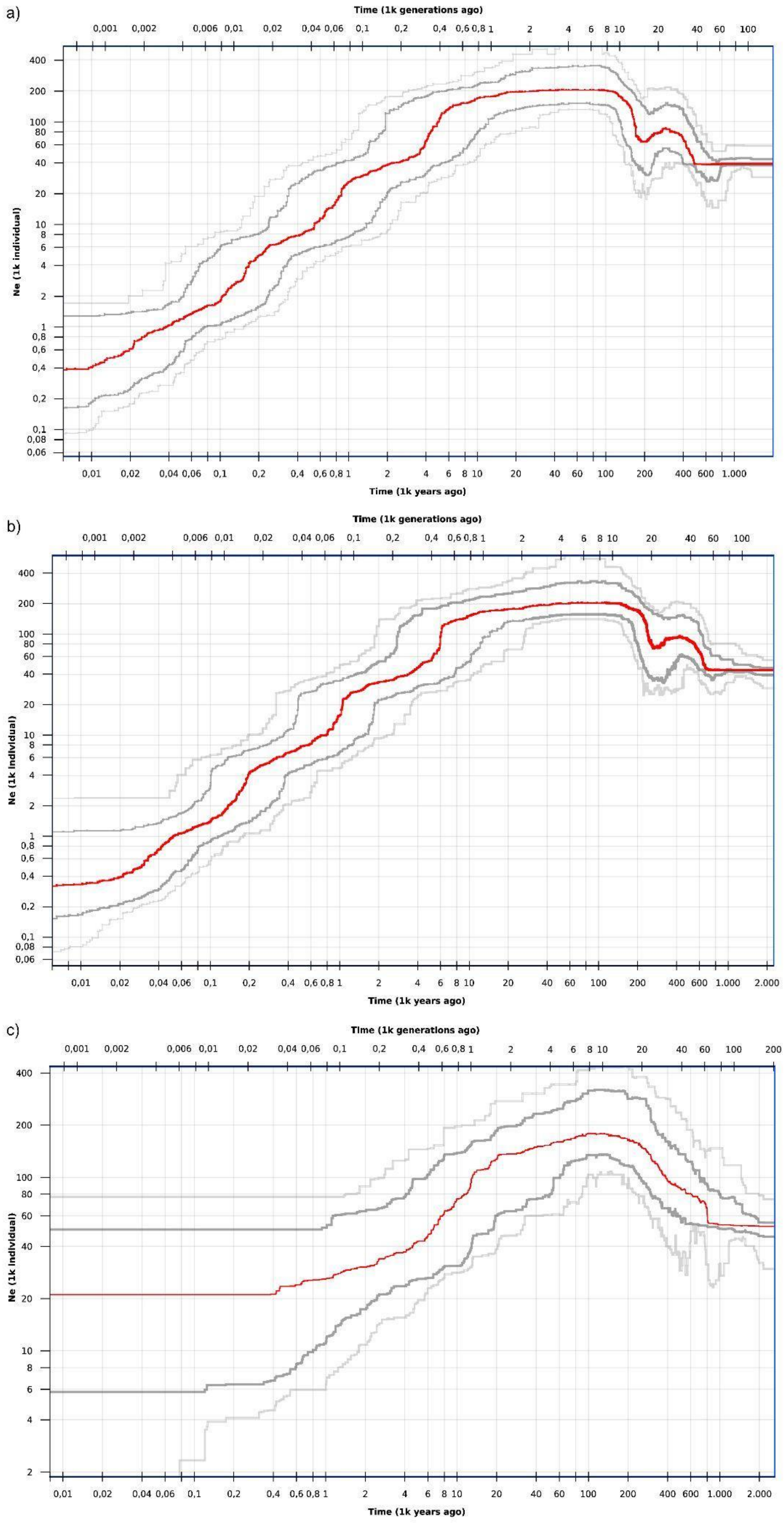
a)
Time (1k generations ago)
Ne (1k individual)
Time (1k years ago)
b)
Time (1k generations ago)
Ne (1k individual)
Time (1k years ago)
c)
Time (1k generations ago)
Ne (1k individual)
Time (1k years ago)

**Supplementary Fig. 17:** Demographic histories of Mediterranean *Puffinus* shearwaters throughout the last million years inferred with StairwayPlot2[9]. Plots for: Balearic shearwaters ($n$ = 20) **(a)**; a joint dataset of all Mediterranean *Puffinus* ($n$ = 32) **(b)**; and Yelkouan shearwaters ($n$ = 12) **(c).** In all cases, lines show the median (red), 75% confidence intervals (dark grey), and 95% confidence intervals (dark grey).

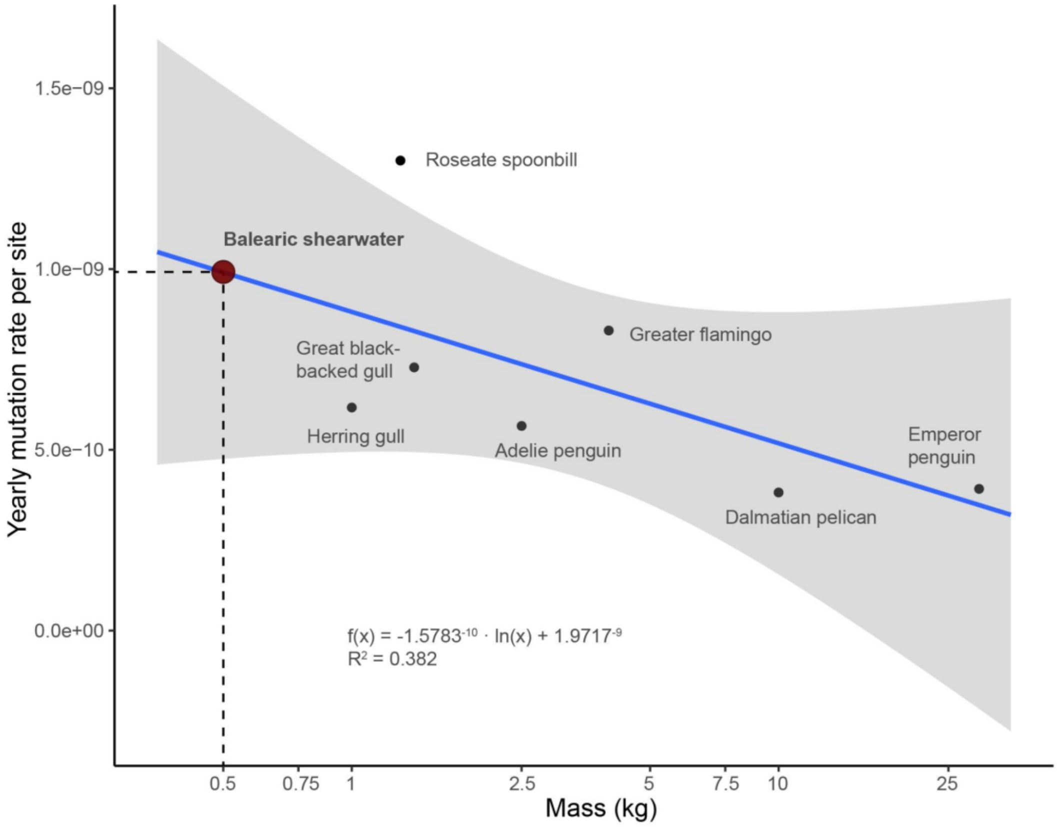


**Supplementary Fig. 18:** Logarithmic regression between body mass and germline mutation rate for the core waterbird species included in Bergeron *et al* (2023)[10]. The intercept of Balearic shearwater, assuming a mean weight of 500g, is shown with dotted lines.

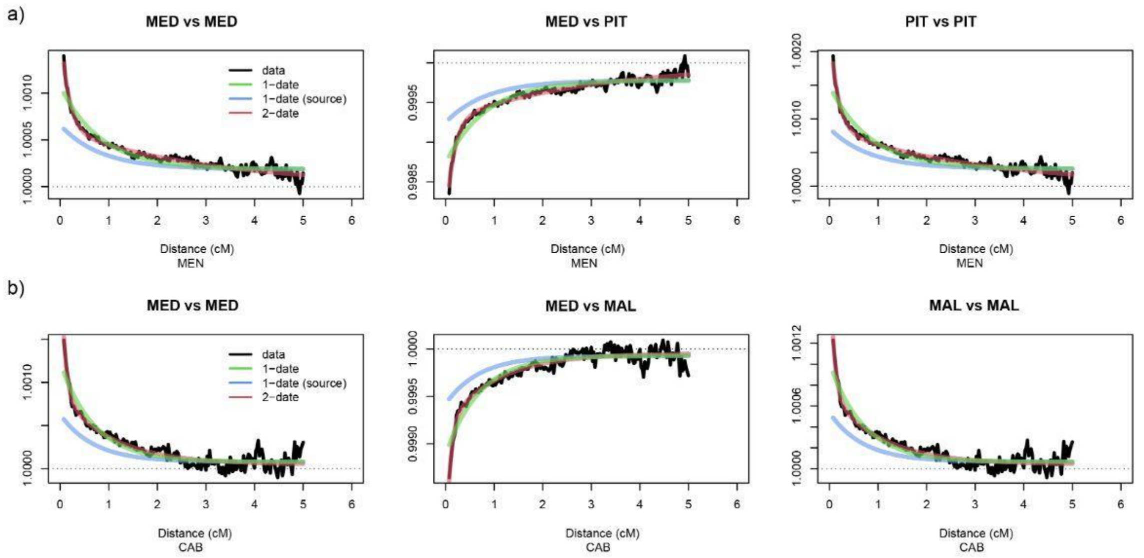


**Supplementary Fig. 19:** Coancestry curves of the minor and major mixing sources inferred in the fastGLOBETROTTER[11] analysis of MEN **(a)** and CAB **(b)**. The y axis gives the weighted probability that haplotypes separated by x distance (as shown in the x axis) belong to different populations. Recent events of admixture will generate long haplotypes for each donor population that are not broken down through recombination; thus, recent admixture will result in less steep curves than ancient admixture. The black points represent the scaled probabilities, while the GLOBETROTTER fitted model is in green (one-date) and red (multiple-dates).

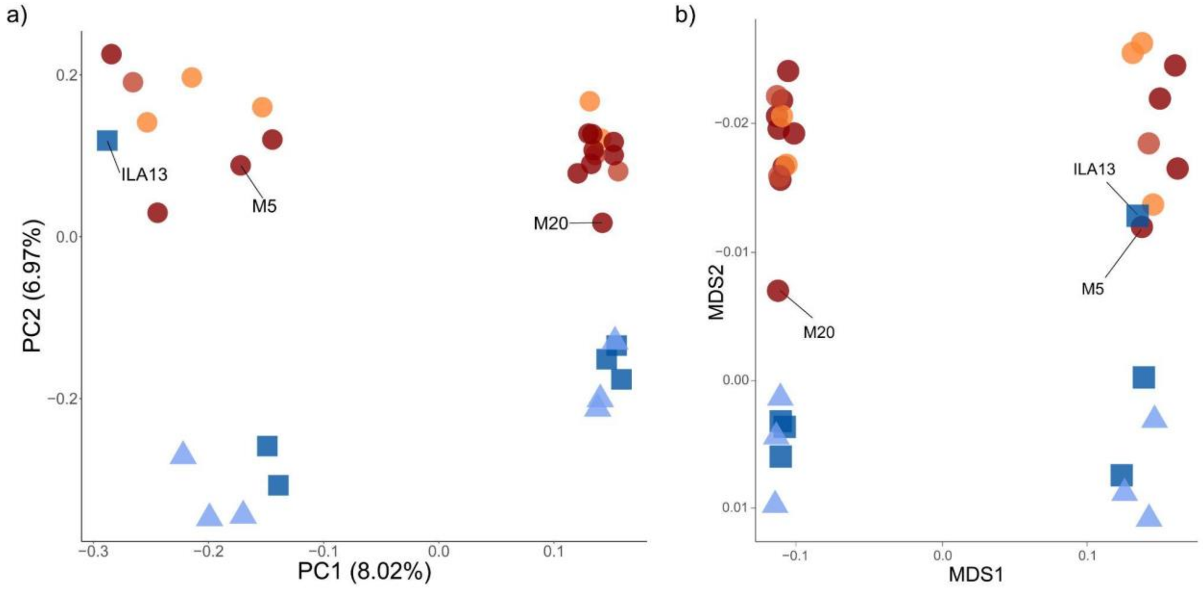


**Supplementary Fig. 20:** PCA **(a)** and multidimensional scaling (MDS) **(b)** based on f4 distances[1] between the studied individuals of Mediterranean *Puffinus* based on 410,281 sex-linked SNPs. Populations are colour-coded according to the legend used in Fig. 1. Labelled individuals were not included in either of the two main clusters (Balearic or Yelkouan shearwaters) in posterior analyses of chromosome Z (e.g., genome scans).

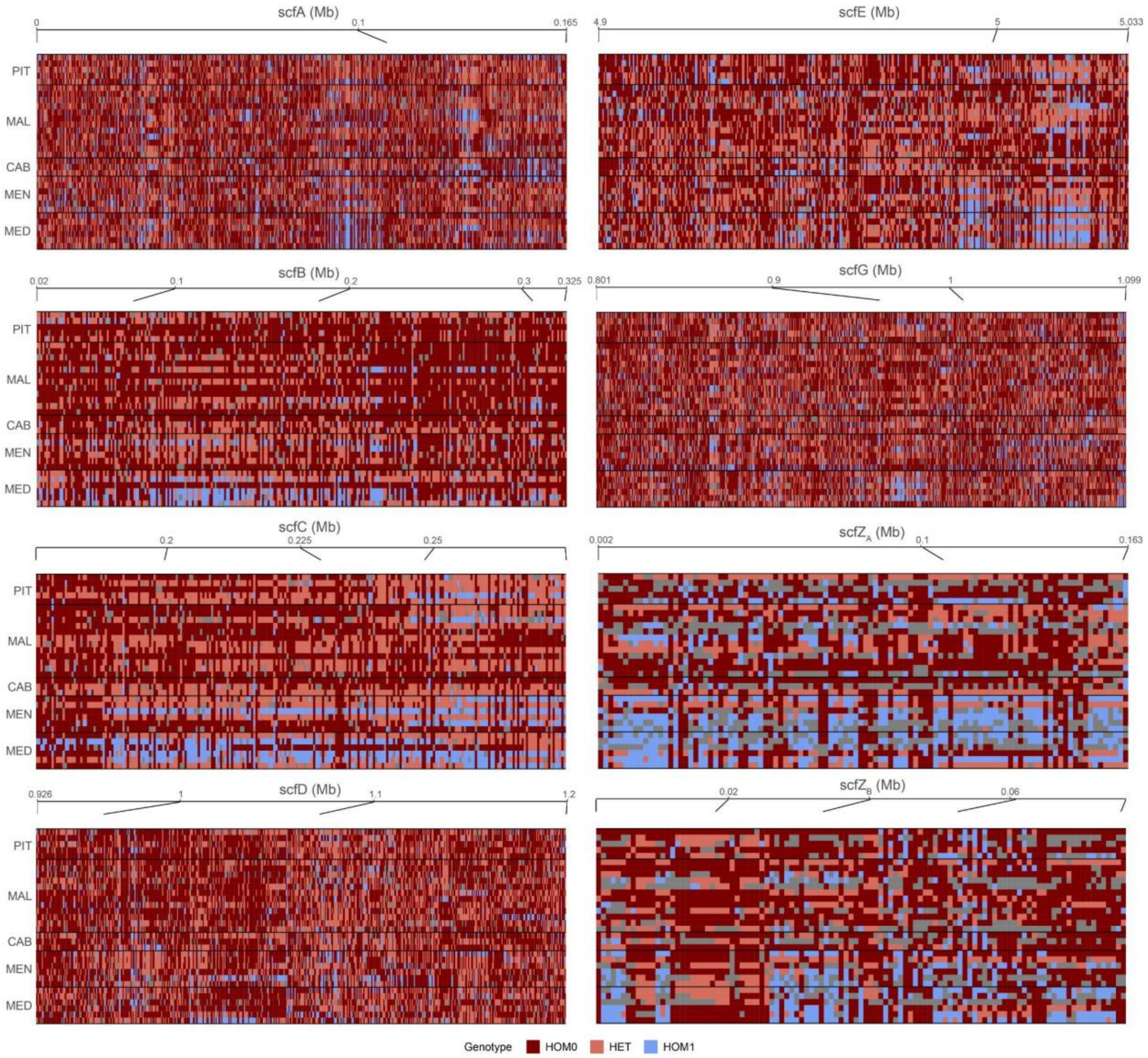


**Supplementary Fig. 21:** Genotype plot of all candidate autosomal windows highlighted by the joint use of $F_{ST}$ and XP-EHH (Fig. 4a). The genotype of each SNP is polarised for the major allele in MAL and depicted as heterozygous (HET: light red), homozygous reference (HOM0: dark red) or homozygous alternative (HOM1: light blue).

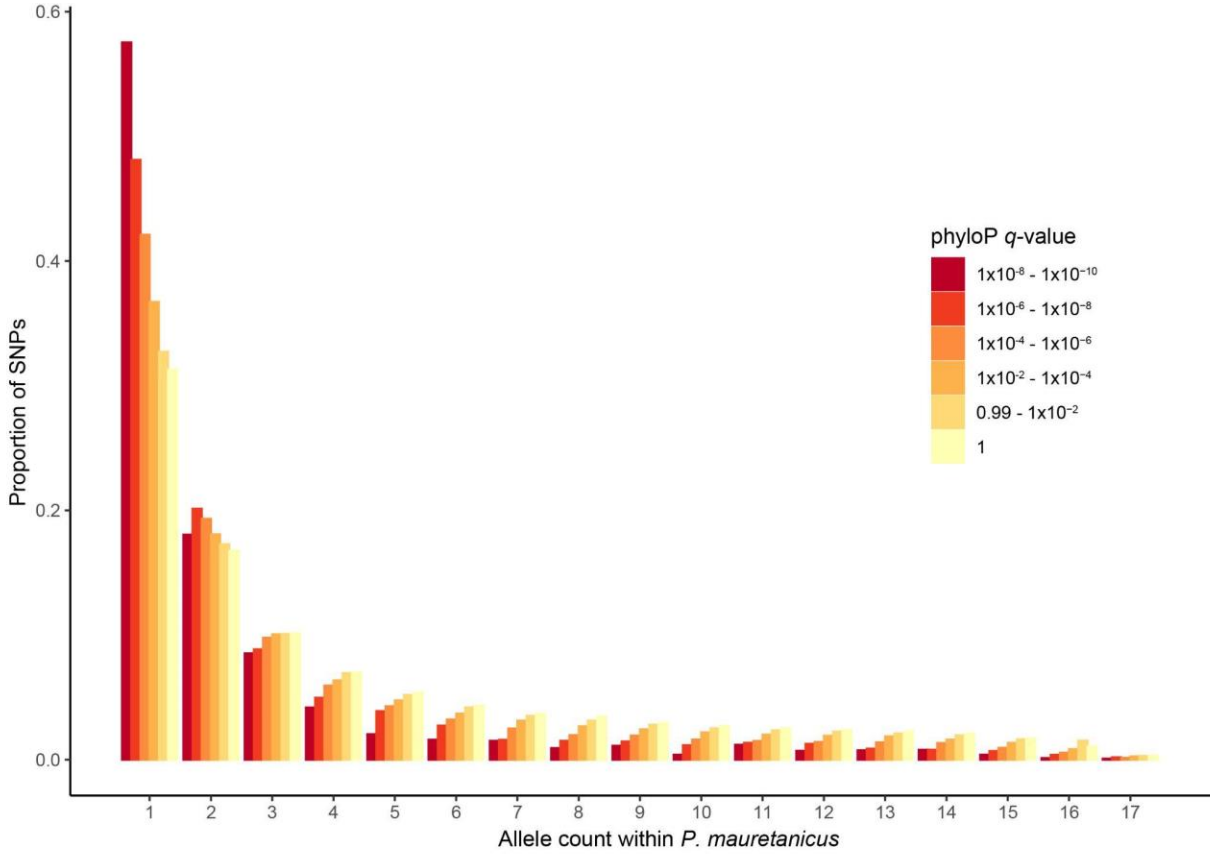


**Supplementary Fig. 22:** Folded site frequency spectrum (SFS) of Balearic shearwater (PIT and MAL) using six SNP datasets containing positions with increasing evolutionary conservation scores - that is, decreasing phyloP[6] *q*-values.

a)

| | N1 | N2 | N3 | N4 | N5 | $N6_B$ | $N6_{NB}$ |
|---|---|---|---|---|---|---|---|
| N1 | 0 | 0 | 0 | 0 | 0 | f*S1*x | 0 |
| N2 | S1 | 0 | 0 | 0 | 0 | 0 | 0 |
| N3 | 0 | $S2*(1-r_3)$ | 0 | 0 | 0 | 0 | 0 |
| N4 | 0 | 0 | $S2*(1-r_4)$ | 0 | 0 | 0 | 0 |
| N5 | 0 | 0 | 0 | $S2*(1-r_5)$ | 0 | 0 | 0 |
| $N6_B$ | 0 | $S2*r_3$ | $S2*r_4$ | $S2*r_5$ | S2 | S3*(1-P) | S2*(1-P) |
| $N6_{NB}$ | 0 | 0 | 0 | 0 | 0 | S3*P | S2*P |

b)

| | N1 | N2 | N3 | N4 | N5 | $N6_B$ |
|---|---|---|---|---|---|---|
| N1 | 0 | 0 | $f'*r_3*S1*x$ | $f'*r_4*S1*x$ | $f'*r_5*S1*x$ | f'*S1*x |
| N2 | S1 | 0 | 0 | 0 | 0 | 0 |
| N3 | 0 | S2 | 0 | 0 | 0 | 0 |
| N4 | 0 | 0 | S3 | 0 | 0 | 0 |
| N5 | 0 | 0 | 0 | S3 | 0 | 0 |
| $N6_B$ | 0 | 0 | 0 | 0 | S3 | S3 |

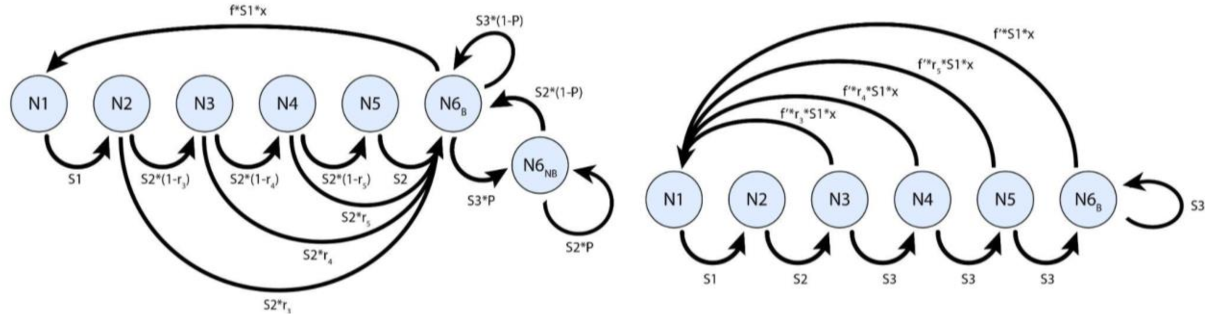


**Supplementary Fig. 23:** Population matrix models and respective life cycle diagrams used to project the Balearic shearwater population under the parameters set in: **a)** Genovart *et al.* (2016)[9] and **b)** a simplified model used in SLiM[4] simulations in the current study. Age-stage classes correspond to unrecruited 1-5 year-old birds (N1-N5), breeding adults ($N6_B$) and sabbatical adults that have bred at least once ($N6_{NB}$). Life-history parameters include: survival probabilities for immatures (S1), non-breeders (S2) and breeders (S3); recruitment probabilities for 3-5 year-old birds ($r_3$-$r_5$); fecundity (f), measured as offspring/female/year; hatching sex-ratio (x); and sabbatical probability (P). In **b)** we introduce a weighted fecundity parameter (f') that is calculated as:

$$f' = f * F_B$$

where $F_B$ is the proportion of surviving breeding adults, calculated as the sum of surviving breeders and floaters that become breeders:

$$F_B = S3*(1-P) + P*S2*(1-P)$$

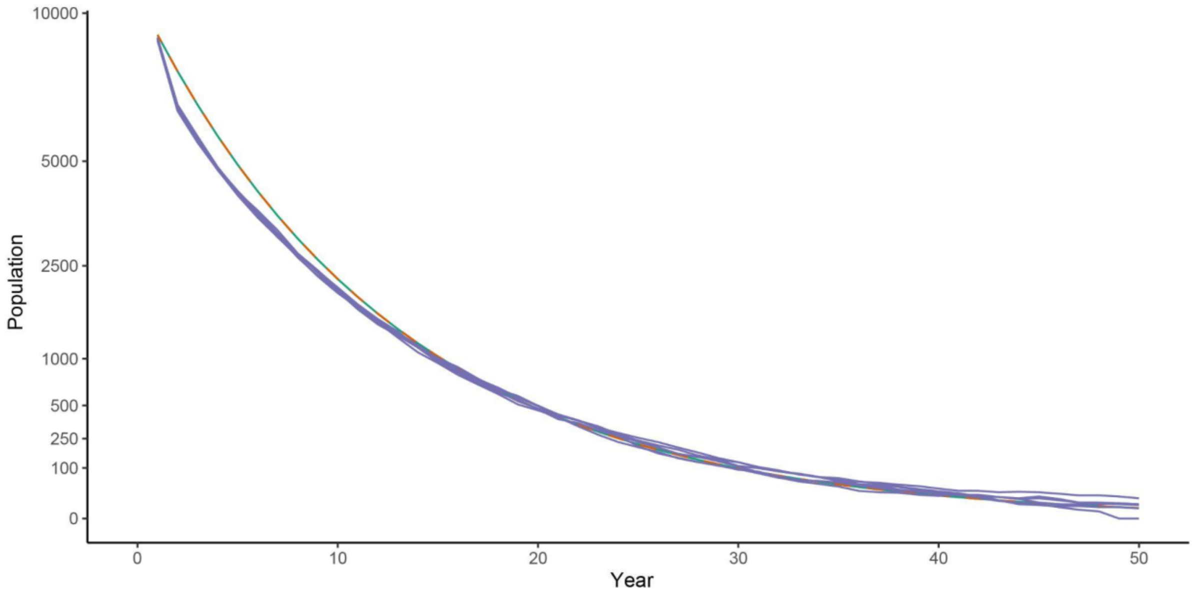


**Supplementary Fig. 24:** Demographic projections for Balearic shearwaters following the survival and reproductive probabilities under current demographic projection. Shown are the deterministic projections using the parameters provided by Genovart *et al.* (2016)[9] (orange); deterministic projections using the simplified Leslie matrix presented in this study (green), and the results of the five SLiM[4] simulations conducted in this study (purple). Notice that deterministic projections using the parameters specified in Genovart *et al.* (2016)[9] and the ones used in this study overlap almost completely, and thus are difficult to discern in this plot.
